# A flexible end-to-end automated sample preparation workflow enables reproducible large-scale bottom-up proteomics

**DOI:** 10.1101/2025.05.23.655690

**Authors:** Sandra Schär, Luca Räss, Liliana Malinovska, Simonas Savickas, Francesca Cavallo, Christopher Below, Marco Tognetti, Polina Shichkova, Benoit Gourdet, Gonzalo Robles, Leo Iu, Jakob Vowinkel, Yuehan Feng, Roland Hjerpe, Roland Bruderer, Lukas Reiter

## Abstract

Bottom-up proteomics holds significant promise for clinical applications due to its high sensitivity and precision, but is limited by labor-intensive, low-throughput sample preparation methods. Advanced automation is essential to enhance throughput, reproducibility, and accuracy and to allow standardization to make bottom-up proteomics amenable for large-scale studies. We developed a fully integrated, automated sample preparation platform that covers the entire process from biological sample input to mass spectrometry-ready peptide output and can be applied on a multitude of biological samples. With this end-to-end solution, we achieved high intra- and inter-plate reproducibility, as well as longitudinal consistency, resulting in precise and reproducible workflows. We showed that our automated workflow surpasses established manual and semi-automated workflows, while improving time efficiency. Finally, we demonstrated the suitability of our automated sample preparation platform for drug development by performing a high-content compound characterization for targeted protein degradation, where high throughput and quantitative accuracy are indispensable. For this, we coupled application-specific workflows to perform proteome profiling and confirm target degradation by precise protein quantification. Overall, our results highlight the selective degradation of specific proteins of interest for ten selected compounds across two cell lines. Thus, the automated sample preparation platform facilitates rapid adaptation to emerging developments in proteomics sample preparation, combining standardization, flexibility, and high-throughput capabilities to drive significant advancements in clinical assays and proteomics research.

## Introduction

High-throughput technologies, including genomics, proteomics, and metabolomics – collectively known as "omics" – have transformed medical research through their ability to provide detailed biological insights into human health and diseases in an efficient and timely manner. Modern genomic platforms enable large-scale projects encompassing several hundreds of samples through automated, high-throughput processes to generate comprehensive datasets and to facilitate rapid and efficient data collection and analysis ^1^. However, this level of advancement has not been uniformly achieved across all omics fields. While proteomics has largely progressed in high-throughput data acquisition, the persistent challenge of labor-intensive and low-throughput sample preparation continues to limit its full potential ^2,3^.

Bottom-up proteomics can provide a comprehensive understanding of molecular mechanisms underlying human diseases by studying proteins. It offers detailed insights into protein abundance, protein-protein interactions, post-translational modifications (PTMs), spatial and temporal profiles, and structural information ^4,5^. Each of these aspects requires specialized workflows for sample preparation, data acquisition, and analysis. Although automation has been introduced for specific steps in sample preparation ^6,7^, a significant challenge remains in developing integrated, end-to-end automated solutions that can accommodate the diverse workflows necessary for comprehensive proteomic analysis ^8^. This challenge is further compounded by the complexity of processing various sample types, such as tissues and body fluids, which demand adaptable and automated processing within a unified system ^9,10^.

Ultimately, to translate bottom-up proteomics into impactful clinical applications, it is essential to meet stringent demands ^8,11^. Process standardization, defined as independence of sample preparation from time, location and operator, is essential to ensure longitudinal reproducibility with high levels of robustness, accuracy and precision. The ability to analyze large number of samples with high consistency is substantial to achieve sufficient statistical power in patient cohorts. In addition, certain sample types are characterized by limited availability and sample fallouts can be detrimental, so low error rates are crucial. Finally, the application in clinical settings requires simplified workflows accessible to personnel without specialized proteomics expertise, rapid turnaround times for clinically relevant insights, and cost-effective operation. Automation of the sample preparation process emerges as a key solution to address those requirements.

In this study, we introduce an end-to-end automated sample preparation platform designed to enhance the performance of bottom-up proteomics and make it suitable for routine large-scale approaches. Targeting throughput ranges of up to 200 samples per day, the platform minimized hands-on time and increased throughput through parallelized workflows. We addressed the requirements for a wide range of workflows and sample types with a modular design that allows the combination of various pre- and post-processing steps for mid- to bulk-ranged sample preparation without compromising data quality. By optimizing liquid handling and processing steps throughout the workflow, we improved processing robustness and minimized sample fallouts. The platform demonstrated high reproducibility, facilitated by processing standardizations such as sample amount normalization, leading to longitudinal consistency over several weeks. It surpassed established workflows in accuracy, precision, and reproducibility, ensuring consistent peptide identification and quantification. Combined with the ability of automated documentation, the platform provides an excellent approach for drug development and biological studies.

## Results

### Implementation of a fully automated workflow provides an integrated end-to-end solution for bottom-up proteomics sample preparation

Large-scale bottom-up proteomic approaches can provide valuable benefits for clinical applications, but require workflows that allow increased throughput, standardization and cost-effectiveness. To meet these requirements, we have developed a fully integrated, automated sample preparation platform that seamlessly integrates with MS acquisition, data processing and downstream analysis (Fig. 1). This platform encompasses a comprehensive workflow that covers the entire process, from biological sample input to mass spectrometry (MS)-ready peptide output. For this, we chose a customized workcell that integrates an 8-channel Hamilton Robotics Microlab STAR M liquid handling robot with a central unit and several third-party devices (Supplementary Fig. 1). This workcell was able to manage all essential tasks in an extensive workflow, as it harbored multiple plate positions with options for both cooling and heating as well as accommodating volumes from nanoliter to milliliter range (Supplementary Fig. 2). We incorporated a positive-pressure module within the central unit for applications requiring positive pressure such as solid-phase extraction or evaporation. In addition, we employed a magnet to interact with magnetic particles, facilitating separation and isolation steps. To regulate temperature-sensitive procedures, we integrated two thermoshakers into the platform, allowing for controlled shaking and/or incubation steps. The integrated robotic arm facilitates seamless interfacing between the on-deck devices and the third-party devices. These include a focused-ultrasonicator for cell or tissue disruption and lysis; a UV-Vis spectrophotometer for concentration determination assays; a plate sealer and peeler for plate coverage and a plate hotel serving as storage for consumables.

**Figure 1.**
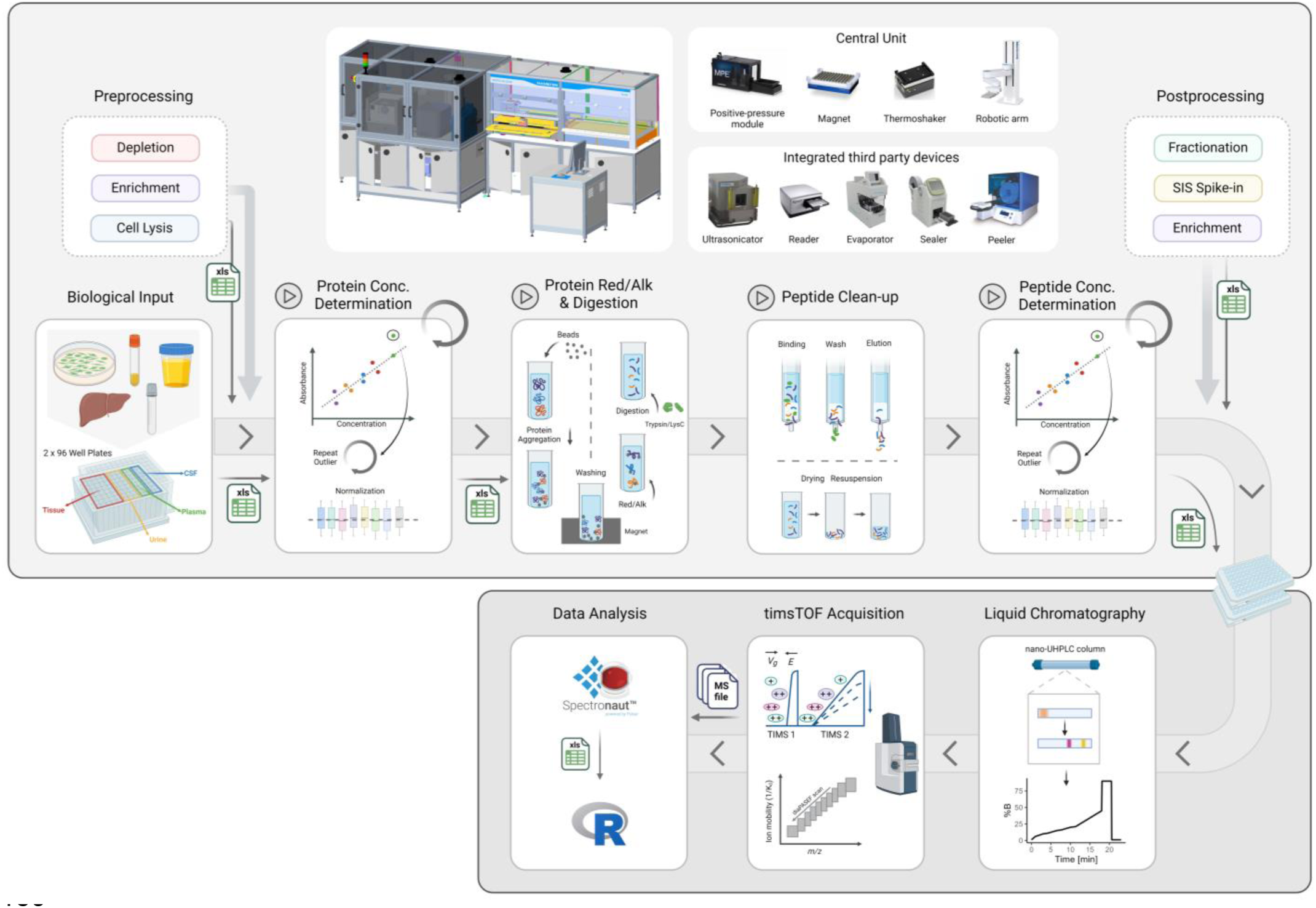
Automated proteomics sample preparation workflow coupled to short-gradient LC-MS acquisition. Schematic overview of the automated sample preparation platform showing the workflow from biological sample input to MS-ready peptide output (upper panel) and subsequent MS acquisition, data processing, and downstream analysis (lower panel). The upper panel shows the customized workcell that includes an 8-channel Hamilton Robotics Microlab STAR M liquid handling robot containing a positive pressure extraction module, 96-well magnet plate, thermoshakers and a robotic SCARA arm. Integrated third-party devices include an ultrasonicator, a microplate spectrophotometer, a nitrogen blowdown evaporator, a plate sealer and a plate peeler. The sample processing workflow is represented in several subpanels where the pre- and post-processing steps are depicted with dashed borders whereas the main workflow is depicted with solid lines. The main workflow starts with protein concentration determination with subsequent sample concentration normalization, followed by sample washes, protein reduction, alkylation, and digestion using the protein aggregation capture process. Peptides are then cleaned using solid-phase extraction, and their concentrations are measured and normalized. The lower panel depicts a 17-minute gradient on a nano ultra-high pressure liquid chromatography (nano-UHPLC) column, utilizing a non-linear gradient. This process is followed by data-independent acquisition parallel accumulation– serial fragmentation (diaPASEF) on the timsTOF HT. The mass spectrometry (MS) raw files generated are subsequently analysed with Spectronaut, and further downstream analysis is performed using R. This figure was generated using BioRender.com.

We designed a workflow that is capable of efficiently processing up to two 96-well plates in a single run. To enhance efficiency, we parallelized the workflow and coordinated it with a dynamic scheduler, distributing tasks across multiple processing units based on real-time workload and resource availability (Supplementary Fig. 3). This approach optimized resource allocation, reduced processing time, and improved overall system performance.

We organized the sample preparation process into three processing blocks: pre-processing, main sample processing, and post-processing (Fig. 1). Pre-processing encompasses preliminary sample-specific steps that samples may undergo prior to being subjected to the main processing. These include cell or tissue disruption, protein depletion and protein enrichment. The main sample processing workflow comprises all steps from the biological sample input leading to MS-ready peptide output. Post-processing covers optional downstream steps such as peptide fractionation, peptide enrichment or spike-in of heavy-labelled peptides for absolute quantification. The sample preparation platform operates on a modular, block wise design, allowing for the desired workflow to be customized from a selection of pre-processing, main and post-processing methods, which can be seamlessly integrated. With this design we ensured smooth assimilation of labware and continuous adaptation of data streams between distinct processing blocks. Moreover, we incorporated various lysis protocols in the pre-processing steps, which enabled the platform to handle a wide range of input materials, including cultured cells, biofluids, and fresh frozen tissue. Overall, this highlights the versatility of the sample preparation platform, which can be optimally tailored to study the various aspects of protein biology in multiple sample types using high-throughput bottom-up proteomics.

We designed the main workflow to handle a predefined input protein amount independent of the sample type. The protein concentration is either assumed based on prior knowledge in the case of well-characterized biofluids such as plasma, CSF and urine, or needs to be determined in the case of cell or tissue lysates (Fig. 1). Therefore, the main method provided the user with the option to either perform or skip the protein quantification assay, based on the sample type. We made sure that if a sample’s concentration surpassed the upper limit of the calibration curve during protein concentration determination, thereby qualifying as an outlier, a reiteration of the assay was triggered automatically, subsequently followed by further dilution of the sample. Next, we applied the protein aggregation capture (PAC) protocol ^12^ on the samples (Fig. 1). In the PAC protocol, aggregated and precipitated proteins are non-specifically immobilized onto microparticles. The immobilized proteins are subjected to washing steps to remove any contaminants, facilitating the retrieval of proteins from a complex background matrix. The proteins undergo reduction, alkylation, and subsequent digestion directly on the beads. The enzymatically digested peptides are acidified and purified via solid-phase extraction. The eluted peptides are dried using a nitrogen blowdown evaporator. Peptides are resuspended, their concentration is determined and then adjusted to a user-defined target concentration, transferred into liquid chromatography coupled to mass spectrometry (LC-MS) compatible 96-well plates and stored at a cooled deck position until further usage (Fig. 1).

With this workflow design, we first tested the hardware functionality and scripting logic and subsequently adjusted the workflow on multiple levels. For this, we focused on reagent-specific liquid handling optimization to account for liquid properties and environmental conditions, achieving accurate and precise pipetting. We minimized dead volumes to conserve expensive and precious reagents, ensuring cost-effectiveness and resource efficiency. Running the protocol repeatedly, we continuously improved process security by advancing error handling and implementing automated error recoveries. This reduced the risk of process failures, eliminated the sample fallout rate and ensured consistent, high-quality outcomes. Finally, we improved the user experience by implementing clear loading instructions, reducing user hands-on time and simplifying the process for end users.

In conclusion, we have developed a highly versatile, multipurpose sample preparation platform. This comprehensive end-to-end automated solution is seamlessly integrated with MS acquisition, data processing and downstream analysis. The modular system minimizes hands-on time while offering high throughput in bottom-up proteomics sample preparation through parallelized work streams.

### Assessment of the automated sample preparation workflow reveals high intra- and interplate as well as longitudinal reproducibility coupled with excellent sensitivity and precision

One key aspect of an automated sample processing workflow is to provide a high level of reproducibility, accuracy and precision. Therefore, we investigated the variability of one sample preparation run. For this, we used HeLa cells as input material and coupled the cell lysis as pre-processing step to the main method. To assess both intra- and interplate reproducibility, 24 replicates of 100 µg protein input were prepared and distributed across two different plates, with each column containing one replicate (Fig. 2A). We analysed the samples using LC-MS and observed highly consistent protein group and peptide identifications across all samples (Supplementary Fig. 4) both within and across plates (Median spearman correlation coefficient ρ = 0.9989 [95% CI: 0.997, 0.999]) (Fig. 2B).

**Figure 2.**
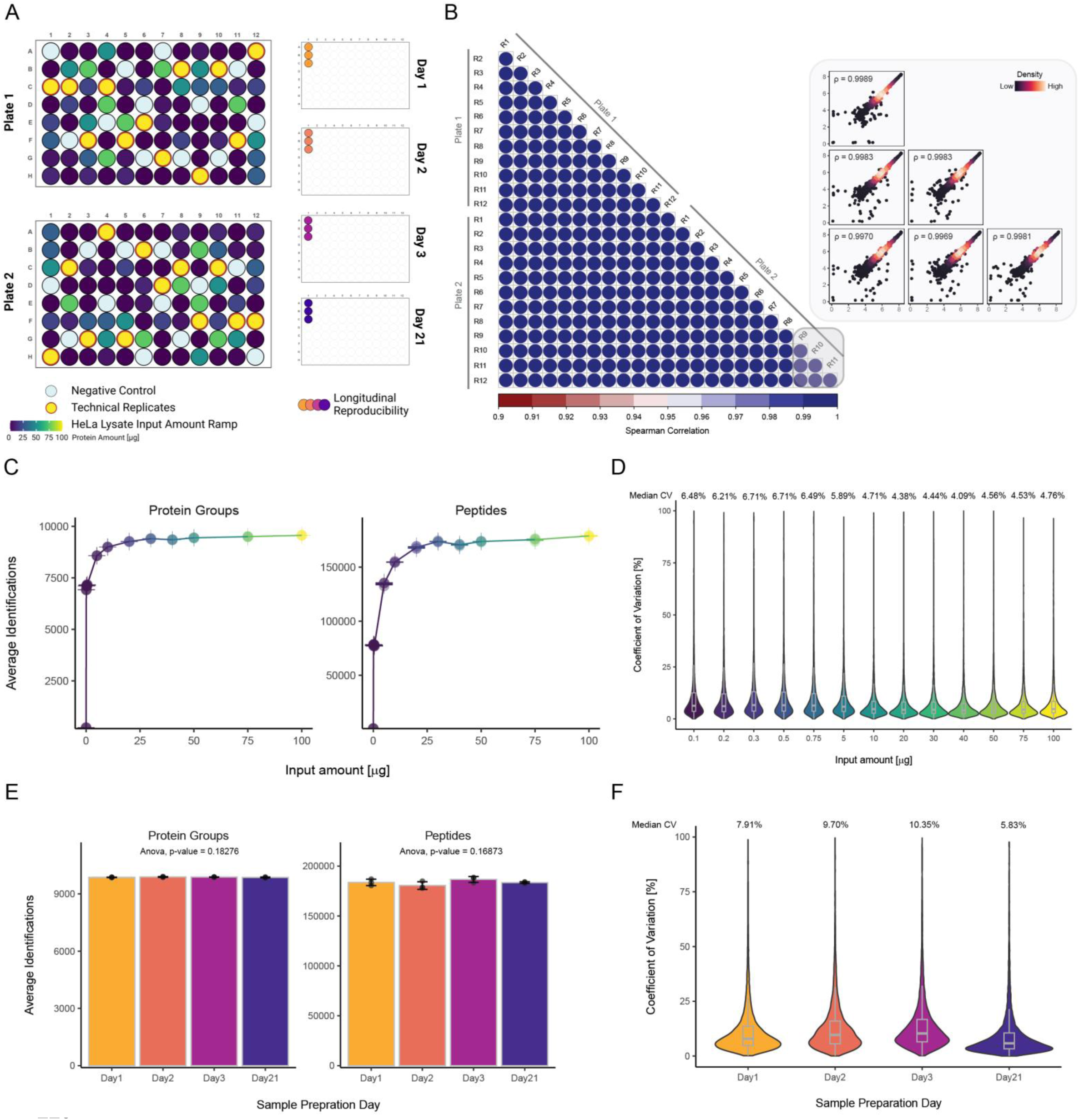
Assessment of reproducibility, sensitivity and precision of the automated sample preparation workflow. **A)** Schematic overview of the experimental design used to evaluate the performance of the automated sample preparation platform, with HeLa cell pellets as input material. Intra- and interplate reproducibility was evaluated by preparing 24 replicates of 100 µg protein and distributing them across two plates with one replicate in each column (yellow dots with red circles). Sensitivity was assessed using a protein input gradient ranging from 0.1 µg to 100 µg (color gradient: dark purple to yellow), including blank replicates (light blue). Longitudinal performance was assessed by processing HeLa cell pellets on four separate days over the course of a month (day 1: light orange, day 2: orange, day 3: dark pink, and day 21: purple). **B)** Spearman correlation heatmap of replicates across two plates (n = 24) with color gradients representing Spearman correlation values ranging from 0.95 (red) to 1.00 (blue) (median spearman correlation coefficient ρ = 0.9989 [95% CI: 0.997, 0.999]). The zoomed-in section highlights representative pairwise scatterplots of protein intensities for three replicates from one plate. Point density is displayed as color gradient. **C)** Number of identified protein groups (left) and peptides (right) across the protein input ramp including negative controls (n = 4, error bars: standard deviation), colored as described in A. **D)** Coefficient of variation (CV) of protein group quantities from the input amount ramp (n = 6) displayed as violin plots, colored as described in A. Embedded box plots display the median (thick lines), interquartile range (25th and 75th percentiles), and whiskers extending to ± (1.58 × interquartile range). Median CVs are displayed above violin plots. **E)** Number of identified protein groups (left) and peptides (right) across samples prepared on days 1, 2, 3, and 21 (n = 3, error bars: standard deviation), colored as described in A. P-values: one-way ANOVA with Tukey’s HSD. **F)** CV of protein group quantities from the longitudinal sample preparation assessment (n = 3) displayed as violin plots with embedded box plots as described in D, colored as described in A. Median CVs are displayed above violin plots. Panel A was generated using BioRender.com.

We tested the occurrence of cross-contamination during the samples processing by measuring 24 negative controls (blanks) across two plates (Fig. 2A). The average of four negative control injections resulted in 2.33% of protein group identifications (220 protein groups) and 0.41% of peptide identifications (709 peptides) compared to conventional injections, indicating minimal cross-contamination relative to sample-containing injections (Fig. 2C). We further tested the stability of the automated sample preparation platform across a large range of protein input amounts. Protein amounts from 0.1 µg up to 100 µg were processed in quadruplicates, distributed across two plates in a single run (Fig. 2A). To avoid overloading the analytical column we normalized the injection amounts (see Methods). All samples with similar injection amount consistently identified between 9’263 and 9’563 protein groups and between 168’276 and 178’878 peptides, indicative of a high similarity in sample recovery, independent of the sample input amount (Fig. 2C). Sub-microgram protein input amounts are of particular interest as they underscore the capability of advanced proteomics techniques to achieve comprehensive protein group identification from minimal sample amounts. Remarkably, while we didn’t optimize the sample preparation for low input, with 0.1 µg protein input, we identified almost 7’000 protein groups, demonstrating the workflow’s exceptional sensitivity. Overall, the increase in protein input correlated with an increase in protein group and peptide identifications but showed consistent and reproducible identifications for each protein input amount. To ensure reliable and consistent protein quantification, we assessed the coefficient of variation (CV) across four replicates for each protein. This evaluation is crucial for determining measurement precision across various protein input amounts. We obtained CVs ranging from 4.09% to 6.49% for each input level, indicating highly consistent protein quantification across replicates (Fig. 2D). This demonstrates efficient and uniform sample preparation from input to MS-ready peptides across a large range of protein input amounts.

We assessed longitudinal performance and inter-day precision by processing HeLa cell lysate in a triplicate on four days (day 1, day 2, day 3 and day 21) i.e., over a time span of three weeks (Fig. 2A). Throughout the analysed timepoints, we consistently identified and quantified 9,573 ± 15 protein groups and 183,542 ± 3,042 peptides (Fig. 2E). We did not observe a statistically significant difference between the timepoints on the protein group level (p-value = 0.18276) or on the peptide level (p-value = 0.16873). We calculated the CV across three samples for each quantified protein, resulting in average CVs of 7.91%, 9.70%, 10.4% and 5.38% for day 1, day 2, day 3 and day 21, respectively. These results reflect the highly consistent protein quantification across replicates of sequentially processed sample plates over the time course of nearly one month (Fig. 2F).

In conclusion, the automated sample preparation allows for the processing of a wide range of protein input amounts without compromising data quality. Furthermore, the implemented workflow demonstrates high intra- and interplate reproducibility, as well as longitudinal consistency over a period of several weeks. This leads to a high standardization in sample preparation and subsequently to precise and reproducible workflows.

### Automated sample preparation workflow achieves better time efficiency and consistent precision in comparison to established methods

The need for reproducible and robust sample preparation for bottom-up proteomics has sparked the development of a multitude of methods over recent years ^10^. Among these, the most commonly used methods are manual in-solution digestion ^13^ and semi-automated protein aggregation capture (PAC) ^12^. In-solution digestion employs a urea-based buffer for protein extraction and denaturation, followed by direct enzymatic digestion within the solution. This method is favored for its simplicity and minimal sample loss. The PAC method is compatible with a wide range of lysis buffers and has been established as semi-automated format, utilizing the magnetic properties of the used beads. For our automated sample preparation platform, we chose the multi-purpose PAC protocol as primary method, as it offers the opportunity to be applied in a fully-automated setting and it is compatible with various lysis settings and effective across diverse biological input materials. We wanted to test the performance of the fully automated sample preparation pipeline and compare it to those of the established workflows regarding efficiency and robustness.

First, we assessed the theoretical processing capacity of all three sample preparation methods. For this, we extrapolated the maximum number of samples processed per working week (SPW) for a workflow including protein concentration determination, protein digestion, peptide clean-up, peptide concentration determination, and finalization. With the robot’s continuous operation, up to 960 SPWs can be processed with only 5 hours of manual intervention throughout the week (Supplementary Fig. 5). This equals 10 96-well plates being processed with the automated platform, whereas the manual and semi-automated workflows can fully process only 3 96-well plates per week. Therefore, the manual and semi-automated workflows are approximately 3.3 times slower and require about six times more manual hands-on time (Supplementary Fig. 5).

Next, we probed the performance of the three sample preparation methods using HEK293 cells as input material (Fig. 3A) For this, we evaluated protein input amounts of 0.1 µg, 1 µg, 10 µg, 50 µg, 75 µg, and 100 µg, processed in sextuplicates for each method (Fig. 3, Supplementary Fig. 6 – 7). Our analysis primarily focused on protein input amounts between 10 to 100 µg, as this range ensures sufficient peptide output for peptide mass normalization in subsequent LC-MS acquisition. We chose 75 µg protein input amount to balance performance, material requirements, and input-output efficiency.

**Figure 3.**
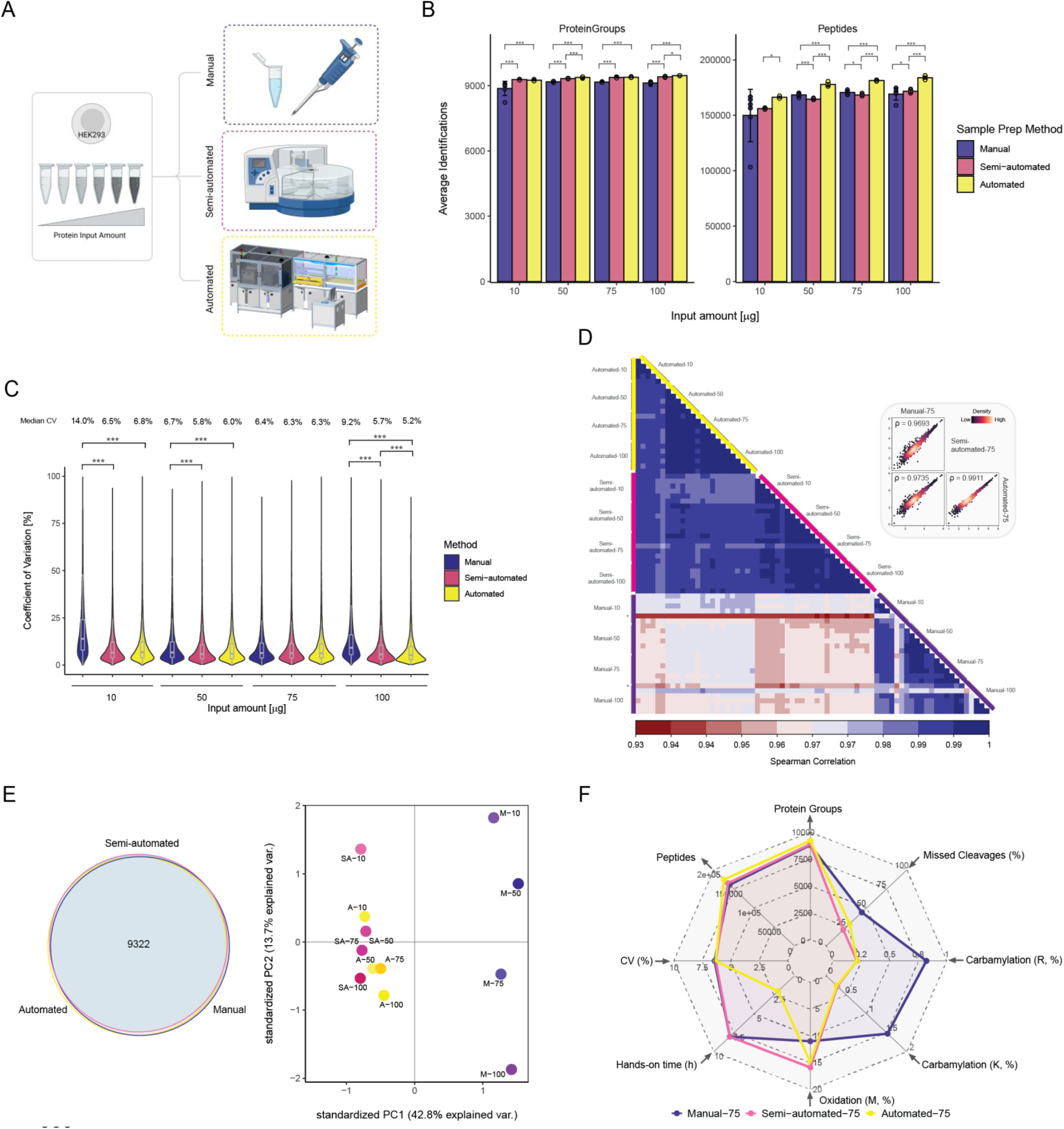
Performance comparison between automated, semi-automated and manual sample processing workflow. **A)** Schematic representation of the experimental design using HEK293 cell pellets for an input amount ramp to compare the automated sample processing (yellow) with a semi-automated (pink) and manual sample preparation workflow (purple). The total processing time (outlined bars) and hands-on time (filled bars) for each workflow are depicted, using the same color scheme as before. Overall processing time (outlined bars) and hands-on time (filled bars) that are required per workflow are depicted with the same coloring as described before. **B)** Number of identified protein groups (left) and peptides (right) across protein input amounts of 10, 50, 75, and 100 µg, prepared using the three workflows (n = 6, error bars: standard deviation). Workflows are colorcoded as described in A. Significance groupings based upon one-way ANOVA with Tukey’s HSD per input amount *p < 0.05, **p < 0.01, ***p < 0.001. **C)** Coefficient of variation (CV) of protein group quantities across the input amount ramp, prepared using the three workflows (n = 6) depicted as violin plots. Workflows are colorcoded as described in A. Embedded box plots display the median (thick lines), interquartile range (25th and 75th percentiles), and whiskers extending to ±1.58 times the interquartile range. Significance groupings based upon one-way ANOVA with Tukey’s HSD per input amount *p < 0.05, **p < 0.01, ***p < 0.001. Median CVs are shown above violin plots. **D)** Spearman correlation heatmap of protein intensities per input amounts across the three workflows (n = 6, workflow types highlighted by colored bars) with color gradients representing Spearman correlation values ranging from 0.93 (red) to 1.00 (blue) (median spearman correlation coefficient ρ = 0.9833 [95% CI: 0.931, 0.997]). Outlier samples are indicated with an asterisk. The zoomed-in view highlights the pairwise scatterplots of protein intensities of one sample with 75 µg input amount from each workflow. Point density is displayed as color gradient. **E)** Venn diagram of protein group overlap (left) and principal component analysis (PCA) based on log2- transformed protein group quantities of different input amounts across the three workflows (right). Work-flows are color-coded as described in A. **F)** Spider web plot visualizing number of protein groups and peptides, protein group CVs, hands-on time, relative proportion of post-translational modifications (e.g., oxidation, carbamylation of lysine (K) and arginine (R)) and missed cleavage ratio for 75 µg protein input amount prepared using the three workflows. Workflows are color-coded as described in A. Panel A was generated using BioRender.com.

First, we compared the number of identified protein groups and peptides (Fig. 3B). The automated workflow showed comparable protein group identifications to the semi-automated PAC workflow, but consistently and significantly outperformed the manual in-solution method (TukeyHSD, p-value < 0.001). Interestingly, the automated workflow also showed a coherently higher identification rate for peptides compared to either, the semi-automated and the manual workflow, for input amounts >10 µg (TukeyHSD, p-value < 0.001). These results emphasize the performance of the automated workflow across various input amounts when compared to two well-established bottom-up proteomics methods. In addition, we calculated the CVs for each quantified protein across the six replicates. The automated sample preparation showed the least variability with median CVs from 5.2% to 6.8%, followed by the semiautomated preparation with median CVs from 5.7% to 6.5%, while the manual preparation had the highest variability with median CVs from 6.4% to 14.0% (Fig. 3C). This demonstrates that the semi-automated as well as the automated workflows exhibit superior reproducibility in comparison to manual preparation, with consistent average CVs across all samples.

To better assess the differences among the methods, we examined the correlation of protein quantities between and within the tested methods (Fig. 3D). Spearman correlation coefficients indicated a very high correlation (>0.97) between the semi-automated and automated sample processing methods. This is likely due to both processes utilizing the same lysis buffer and following the same sample processing principle. The correlation remained high (>0.93) between the automated and manually processed samples, reinforcing the robustness of both procedures. Additionally, we observed an exceptionally high correlation even between different input amounts within each preparation method, further highlighting the consistency and reliability of these methods across varying protein input amounts (median spearman correlation coefficient ρ = 0.9833 [95% CI: 0.931, 0.997]). However, it is important to acknowledge the presence of outliers specifically in the manually prepared samples, which indicates variability in the manual process.

Beyond the number of proteins and the precision of each workflow, it was critical to consider whether the different methods vary in the identity and quantity of the proteins they extract. Therefore, we compared the overlap and uniqueness of protein group identifications across the three sample preparation strategies using 75 µg input amount (Fig. 3E). Notably, 9,322 protein groups, representing 93.4% of the total, were shared across all three workflows. This substantial overlap indicates that the majority of proteins are effectively extracted and processed regardless of the sample preparation protocol. A principal component analysis (PCA) of protein quantifications delineated distinct clusters along the first principal component (PC1), accounting for 42.8% variance. These clusters correspond to different sample preparation methods, indicating method-dependent variability. While the identified proteins were largely consistent across methods, subtle differences in protein quantities were evident corroborating the effect of different biochemistry underlying sample preparation methods.

The main differences among the three sample-preparation methods are summarized in Figure 3F and Supplementary Figure 8. We tested whether individual sample preparation methods are prone to protein modification artifacts. Indeed, we observed method-specific modifications and adducts. For example, peptides originating from samples treated with urea-buffer showed increased levels of carbamylation on arginine and lysine, a well-known modification associated with this buffer compound ^14^. The semi-automated as well as the automated protocol revealed higher oxidation rates compared to the manual protocol. We assume that this is due to increased exposure to air from the additional pipetting and shaking steps throughout the PAC protocol. We compared the incidence of missed cleavages to evaluate the efficiency of enzymatic digestion and found a higher rate in the manual workflow compared to the other two methods. This indicates suboptimal digestion with 44% of peptides having a missed cleavage site.

In summary, the comparison of manual, semi-automated, and automated sample preparation workflows demonstrated the robust performance of all three methods. We showed that the automated platform not only matches but even surpasses established workflows in identification, accuracy, precision reproducibility as well as efficiency.

### Combination of the automated sample preparation platform and high-throughput MS acquisition allows deep biological insights in short time

With a high-performance sample preparation platform capable of processing numerous samples with minimal hands-on time available, we proceeded to integrate it with our established high-throughput LC-MS setup within Biognosys’ TrueDiscovery and TrueSignature platform to pave the way for high-throughput proteomic assays. To demonstrate the suitability of the setup with commonly employed cell line screening approaches, we chose to process a panel of nine widely used cell lines (Fig. 4A). These comprised primarily cancerous and tissue-derived cells: A549 (lung carcinoma), HCT116 (colorectal carcinoma), K562 (chronic myeloid leukemia), MCF7 (mammary carcinoma), NB4 (acute promyelocytic leukemia), SKOV3 (ovarian carcinoma), U251 (glioblastoma), and U937 (histiocytic lymphoma). Additionally, we included the non-malignant cell line HEK293 and patient-derived peripheral blood mononuclear cells (PBMCs) ^15–19^. We performed proteomic profiling across three biological replicates of all samples using the automated sample preparation workflow followed by diaPASEF acquisition on a timsTOF HT instrument. The cumulative processing time for the 30 samples was 31 hours, encompassing both sample preparation and acquisition. Notably, manual handling constituted only 10% of the total duration. In the resulting extensive biological dataset, we discovered a cumulative total of 11’174 unique protein groups and an average of 9’302 protein groups across the complete biological dataset. For individual samples we identified between 8’959 and 9,696 distinct protein groups as well as between 104’563 and 144’877 peptides (Fig. 4B). As expected, the CVs for each protein group quantity calculated across replicates was low, within the range of 5.80% to 9.03% for all samples (Supplementary Fig. 9). This finding underlines the high reproducibility of the experimental approach. Next, we examined the biological differences of the samples using a principal component analysis (PCA) based on protein quantities across cell samples (Fig. 4C). Each distinct cell line was clearly separated in PCA space, with PBMCs showing the greatest distance from the other nine cell line samples. This shows that the cellular heterogeneity in patient-derived PBMCs was preserved, while the cultured cell lines displayed a higher level of relative homogeneity. Nevertheless, the individual cell lines clustered as distinct groups, demonstrating that their individual specific traits are preserved and readily detectable when processed with the automated sample processing platform.

**Figure 4.**
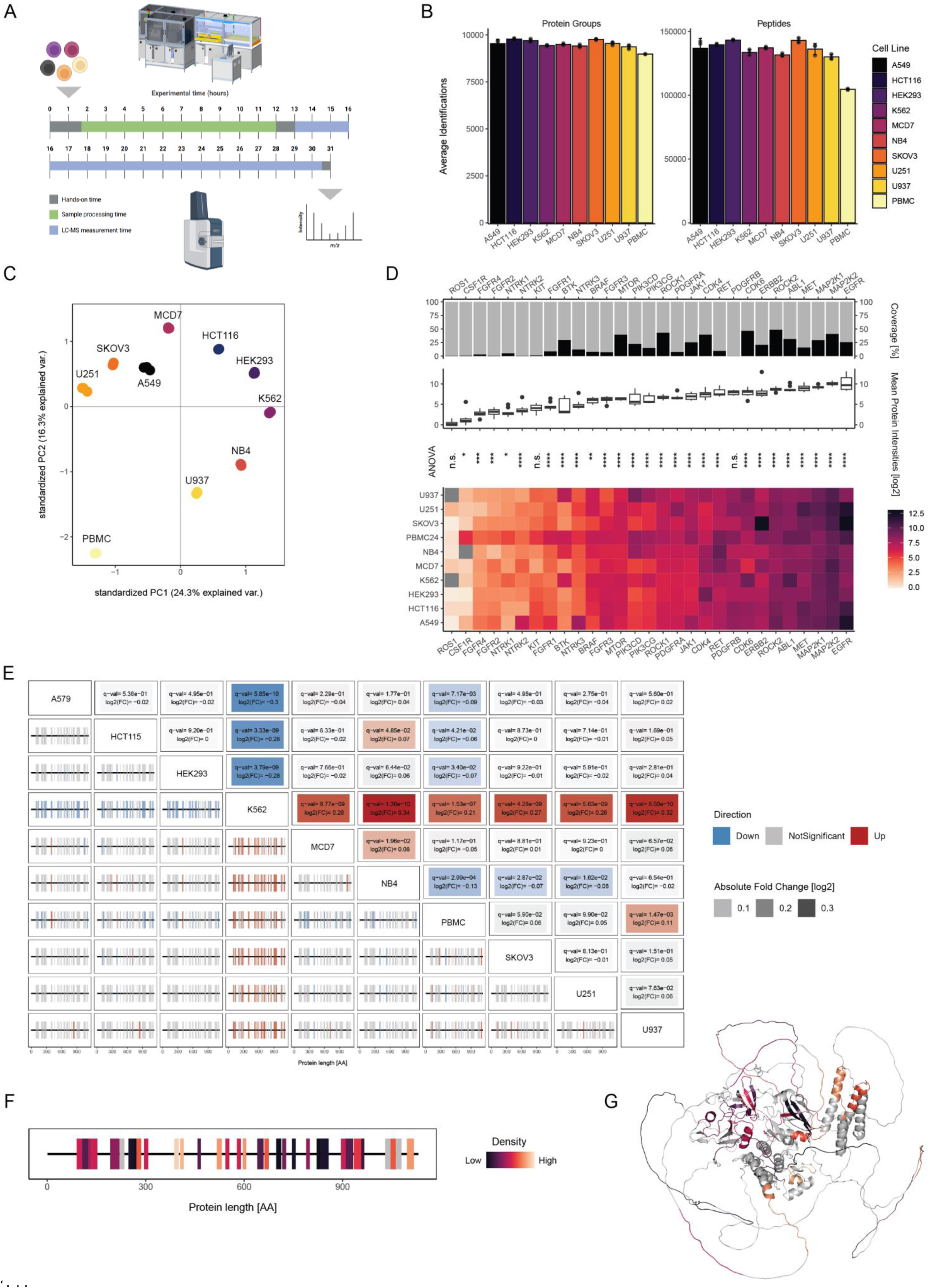
Characterization of multiple cell lines using the automated sample preparation workflow coupled to short-gradient LC-MS acquisition. **A**) Schematic representation of the experimental design using a panel of nine cell lines: A549 (lung carcinoma), HCT116 (colorectal carcinoma), K562 (chronic myeloid leukemia), MCF7 (breast carcinoma), NB4 (acute promyelocytic leukemia), SKOV3 (ovarian carcinoma), U251 (glioblastoma), and U937 (histiocytic lymphoma), HEK293 (human embryonic kidney) and patient-derived peripheral blood mononuclear cells (PBMCs). Samples were prepared using the automated sample preparation workflow, followed by short-gradient LC-MS acquisition. Total sample preparation time was 10 hours (green), acquisition was completed in 17.5 hours (blue) with total hands-on time of 2.5 hours (grey). **B**) Number of identified protein groups (left) and peptides (right) across distinct cell line replicates (error bars: standard deviation). Different cell lines are distinguished using a color gradient ranging from black to light yellow. **C**) Principal component analysis (PCA) of protein expression levels per cell line. Cell line color coded as described in B. **D**) Bar plots illustrating peptide coverage of selected kinases across cell lines (top panel). Box plots showing the median (thick lines), interquartile range (25th and 75th percentiles), and whiskers extending to ±1.58 times the interquartile range for the intensities of selected kinases across cell lines (middle panel). Heatmap displaying the mean protein intensities of selected kinases across cell lines (lower panel). One-way ANOVA tests for differences in mean protein intensities between cell lines. Significance labels derived from Tukey’s HSD * = p < 0.05, ** = p < 0.01, *** = p < 0.001, **** = p < 0.0001, n.s. = not significant. **E**) Upper triangle: Pairwise differential abundance testing between cell lines on protein group level. Log2 fold changes (FC) and adjusted p-values (Benjamini-Hochberg) are displayed. Significant changes in intensity between two distinct cell lines (|log2(FC)| >1, q-value < 0.05) are shown in red (upregulation) or blue (downregulation) depending on the directionality of change. Non-significant changes are displayed in grey. Extent of change (|log2(FC)|) represented in color intensity. Lower triangle: barcodes represent the detected peptides along the ABL proto-oncogene 1 (ABL1) protein from the N to C terminus. Each vertical bar corresponds to a peptide detected in the samples. Peptides with a significant change in intensity are shown in red (upregulation) or blue (downregulation) as described above. Peptides that were detected but did not change between conditions are displayed in grey, while regions not covered are not shown. **F**) Cumulative analysis of all pairwise comparison to K562. Barcode representation as described in E. The coloring of the peptides represents the frequency of a significant change across all comparisons. **G**) Protein structure of ABL1 (AlphaFold prediction: AF-P00519-F1) with mapped frequency of significant peptide changes as described in F. Panel A was generated using BioRender.com.

Some of the most prominent functional differences between different cell types are represented through differential expression profiles of kinases and ubiquitin ligases ^20,21^. Moreover, due to their pivotal role in cell signaling and regulation, kinases and ubiquitin ligases are important targets in cancer drug screening approaches. Thus, we assessed the protein expression pattern of selected kinases ^22–26^ and ligases ^27,28^ in our data (Fig. 4D, Supplementary Fig. 10). We observed significant differences in the expression patterns of most of the kinases and ligases (TukeyHSD, p-value < 0.05), recapitulating the expected functional heterogeneity. The comprehensive data enabled us to further investigate the differential expression patterns of specific proteins. For this, we focused on a protein with an overall high expression level and a high sequence coverage: ABL proto-oncogene 1 (ABL1), a non-receptor tyrosine kinase. ABL1 has been associated with chronic myeloid leukemia, where its abnormal activation leads to uncontrolled cell proliferation and survival ^29^. In agreement with this, we observed the most pronounced expression differences in K562 cells (Fig. 4E), a cell line derived from a chronic myelogenous leukemia patient ^30^. Moreover, we observed a highly consistent peptide quantification across all identified peptides of a given protein (Fig. 4F, G), underscoring the robustness and reproducibility of our workflow. Taken together, these data demonstrate the power of the platform to rapidly generate accurate high content data that can be interrogated to drive decision making in experimental approaches for drug discovery.

### Automated sample preparation coupled to LC-MS can be utilized for target identification, quantification and selectivity profiling of molecular degraders

Targeted Protein Degradation (TPD) is a pharmaceutical modality that enables modulation of proteins traditionally considered undruggable by conventional small molecules ^31–33^. TPD approaches recruit the cellular machinery to drive degradation of pathogenic proteins, most often by employing molecular glue degraders or proteolysis targeting chimeras (PROTACs) (Fig. 5A, top panel). Molecular glues act by promoting or stabilizing protein-protein interactions between the E3 ligase and a protein of interest (POI), leading to selective ubiquitylation and degradation of the target ^34^. PROTACs are heterobifunctional small molecules that recruit an E3 ubiquitin ligase to the POI, facilitating its ubiquitylation and subsequent degradation by the ubiquitin-proteasome system ^35^.

**Figure 5.**
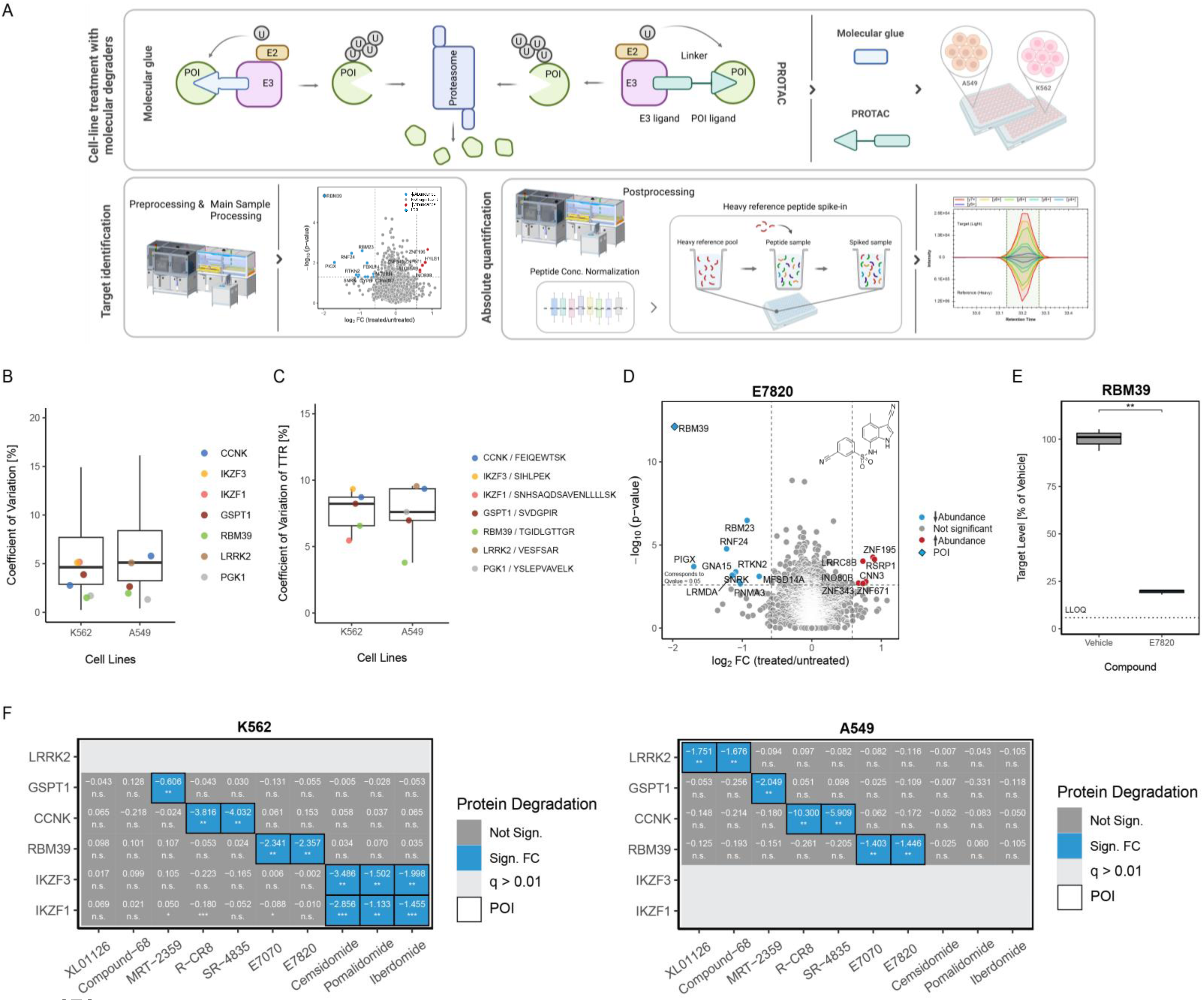
Automated sample preparation coupled to LC-MS can be utilized for target identification and quantification of molecular degraders. **A**) Schematic representation of the experimental design. Proteolysis targeting chimeras (PROTACs) and molecular glues facilitate the binding of E3 ubiquitin ligase to a protein of interest (POI), leading to selective ubiquitylation and subsequent degradation by the ubiquitin-proteasome system. Two cell lines were treated with selected molecular degraders, followed by identification of targets through differential analysis and confirmation of their degradation by measuring protein levels. **B**) Coefficient of variation (CV) of all protein group quantities in five vehicle samples of A549 and K562 cell samples depicted as boxplots displaying the median (thick lines), interquartile range (25th and 75th percentiles), and whiskers extending to ±1.58 times the interquartile range. POIs and control proteins are color-coded. **C**) CV of target-to-reference ratio (TTR) of selected peptides for each POI (color-coded, protein origin and peptide sequence shown in legend) used for quantification in five vehicle samples of A549 and K562 cells depicted as boxplots displaying the median (thick lines), interquartile range (25th and 75th percentiles), and whiskers extending to ±1.58 times the interquartile range. **D**) Volcano plot illustrating the differentially abundant proteins identified through proteome profiling between K562 cell samples (n = 5) treated with the molecular degrader compound E7820 including the corresponding molecular structure. Proteins are color-coded based on their statistical significance and fold change: red indicates increased abundance, blue indicates decreased abundance (|log2(FC)| > 0.58, q-value < 0.05) and grey indicates no significant change (|log2(FC)| ≤ 0.58, q-value > 0.05). E) Boxplot showing the absolute amount of the POIs in K562 cell samples (n = 3) treated with the molecular degrader compound E7820, compared to DMSO-treated samples (vehicle). T-test based p-values are indicated in the plots: * = p < 0.05, ** = p < 0.01, *** = p < 0.001, **** = p < 0.0001, n.s. = not significant. Lower limit of quantification (LLOQ) is indicated as dotted line. **F**) Heatmaps displaying fold changes from absolute quantities in K562 cells (left) and A549 cells (right) results for molecular degrader compounds (x-axis) and identified targets (y-axis). Fold change is color-coded: blue for significantly negative (log2 |FC| ≥ 0.58) and grey for no significant change (log2 |FC| ≤ 0.58). T-test based p-values: * = p < 0.05, ** = p < 0.01, *** = p < 0.001, **** = p < 0.0001, n.s. = not significant. POIs with its selected peptides with a q-value of > 0.01 are greyed out. Anticipated POIs per compound are framed in black. Panel A was generated using BioRender.com.

High-content proteomic profiling for molecular degrader characterization involves the treatment of cells with a collection of compounds, identification of targets through differential expression analysis, and the confirmation of their degradation by quantification of protein levels. This characterization approach can greatly benefit from automated high-throughput platforms (Fig. 5A). We demonstrated this by simultaneously screening two cell lines treated with ten preclinical and clinical-stage molecular degraders: eight molecular glue degraders and two PROTACs, each targeting one of six disease-relevant proteins known to be involved in oncology ^36–41^ or neurodegenerative disorders ^42^ (Supplementary Data 4, Supplementary Fig. 18). Based on the previously measured protein expression levels across different cell lines (Supplementary Fig. 11), we selected two cell lines: the chronic myelogenous leukemia continuous cell line K562 and the lung carcinoma epithelial cell line A549. Additional testing showed that 225,000 cells per well provided sufficient protein and peptide yield for follow-up analysis, stable identification and low variations (for details see Supplementary Information). We treated the two cell lines with either dimethyl sulfoxide (DMSO) as negative control (vehicle) or with each compound at the indicated concentration (Supplementary Data 4) for five hours to ensure an observable effect.

We performed proteomic profiling using the TrueDiscovery platform of Biognosys for each sample to assess degradation of targets of interest. Our analysis revealed robust and reproducible protein and peptide identifications across both cell lines (Supplementary Fig. 15). Additionally, we calculated the CVs for each quantified protein across five vehicle-treated samples (Fig. 5B). The median protein CVs were 4.9% for K562 cell samples and 5.5% for A549 cell samples, which confirms the high reproducibility of the experimental approach. The proteome profiling data can also be used for the identification of suitable peptides for quantitative analysis of POI protein levels ^43^. We used the previously acquired data of the two cell lines and selected peptides using tightly defined criteria (see Methods for details). For the reference-based quantification of the selected POIs using Biognosys’ TrueSignature platform, we used the automated post-processing step to spike-in heavy-labelled peptides into the cell lysates.

We assessed the pipetting precision of the spike-in by calculating the CVs of the target-to-reference (TTR) ratio of peptides in five vehicle-treated samples (Fig. 5C). Stable TTRs are key for reference-based quantification, as they indicate precise pipetting of the reference and thus low technical variations. We observed median peptide CVs of 8.2% in K562 samples and 7.3% in A549 samples (Fig. 5C), which highlights the robustness and reproducibility of the automated workflow, and particularly of the post-processing system.

To monitor protein degradation, we performed differential expression analysis of the proteomic data and assessed proteins that showed a significant change in abundance in compound- vs. vehicle-treated conditions (|log2(FC)|>0.58, q<0.05) (Fig. 5D, Supplementary Fig. 16 – 17). Five out of ten compounds led to the known POI being ranked as the top candidate among the down-regulated proteins in the cell line with the highest POI expression. Figure 5D illustrates the results of K562 cells treated with compound E7820 – its POI RBM39 is the most down-regulated protein (evaluation to other compounds see Supplementary Information). For compounds XL01126 and Compound-68, POIs were identified only in A549 cells due to low expression in K562 cells. Conversely, for Cemsidomide, Pomalidomide, and Iberdomide, POIs were identified only in K562 cells due to low expression in A549 cells (Supplementary Fig. 11). We confirmed the successful degradation of each POI by quantitative analysis of the protein levels (Fig. 5E, Supplementary Fig. 16 – 17). Figure 5E illustrates an 80% degradation of RBM39 protein in K562 cells treated with E782 compared to vehicle-treated cells, indicating a significant reduction in protein levels upon treatment. Across all compounds, we demonstrated 35–95% degradation of the known POI in the cell line with the highest POI expression (Fig. 5E, Supplementary Fig. 16 – 17). Overall, our results highlight the selective degradation of specific POIs for each compound in the two cell lines (Fig. 5F). Interestingly, we observed cell line-dependent differences, particularly with certain POIs, varying from high endogenous abundance in one cell line to inconclusive measurements below limit of quantification (<BLQ) in the other one. This illustrates the critical importance of selecting appropriate cell lines or incorporating multiple cell lines in the characterization approach for TPDs.

In conclusion, we demonstrated that the automated sample preparation workflow is highly suitable for high-content compound characterization due to its robustness, reproducibility and efficiency. This workflow encompasses both target identification and subsequent quantification evaluation, ensuring comprehensive analysis. The automated solution for targeted assays facilitates longitudinal stable reference peptide spike-ins for measuring protein levels, which is crucial for maintaining consistency over time. This makes the workflow particularly well-suited for drug development, where precise and reliable quantification of target proteins is essential.

## Discussion

In this work, we developed an end-to-end automated sample preparation platform designed to enhance the performance of bottom-up proteomics. Its modular, block wise design allows a seamless integration of customizable workflows with various combinations of pre- and post-processing methods. It efficiently processes a wide range of protein input amounts and various sample types simultaneously, without compromising data quality. The implemented workflow demonstrates high reproducibility through longitudinal consistency over several weeks and exhibits high intra- and inter-plate reproducibility. The marked increase in efficiency, reduced error rates and optimized user experience makes it an ideal system for a cost-efficient and simple application that does not require extensive expert knowledge. We showed that the platform is suitable for high-content compound characterization where precise and reliable quantification of target proteins is essential especially for downstream clinical research.

Sample preparation remains one of the most critical and error-prone steps in bottom-up proteomics, since it encompasses multiple sequential processes such as sample preprocessing, protein quantification, reduction and alkylation, enzymatic digestion, peptide quantification, and post-processing. In standard workflows, these steps can involve up to 15 pipetting steps per sample, which translates to approximately 2’880 pipetting steps for a 192-sample batch. Each step introduces a potential source of technical variability, which can accumulate throughout the process. However, with automated liquid handling, pipetting steps can be parameter-controlled and standardized over time. While individual automation solutions for selected steps, such as automated digestion or protein quantification, have been proposed, a comprehensive automation of the full sample preparation pipeline has remained limited ^44–46^. Our end-to-end automated solution addresses this gap by fully integrating all key steps from biological sample input to MS-ready peptide output within a single, autonomous workflow. In contrast to existing systems that often require manual interventions, such as sample dilution when the concentration is outside of the calibration range ^47^, centrifugation steps or manual sealing/unsealing of plates ^7^, our system is designed for hands-free processing with built-in redundancies, automated recovery options, and real-time error handling, ensuring uninterrupted operation. This complete automation minimizes human interaction, reduces the risk of contamination or sample loss, and enhances reproducibility through standardized pipetting across batches and time points. By decoupling the sample preparation process from operator, time, and location variability, our system facilitates robust scaling and seamless integration into regulated or clinical environments, supported by comprehensive logbooks and audit trails.

Recent advancements in LC-MS measurement time have significantly enhanced bottom-up proteomics. Within five years, the time required to identify 10,000 protein groups has been reduced threefold, from 100 to 30 minutes, enabled by advances in MS instrumentation and the adoption of shorter LC gradients ^48,49^. Consequently, to keep pace with the speed of LCMS acquisition, sample preparation must be scaled up accordingly, which can be best achieved through the use of robotics. Our automated platform processed 30 cell samples within 10 hours, requiring only 2.5 hours of manual hands-on time. When scaled to 192 samples, the workflow, enabled by parallelized processing, completes sample preparation from biological input to MS-ready output within 24 hours. Notably, the required manual hands-on time increases marginally to approximately 3 hours, underscoring the efficiency gains achieved through automation and parallelization. Extrapolating this capacity to continuous weekly operation, our system can process approximately 960 samples per work week with an estimated 5 hours of total manual interaction, thereby enabling high-throughput sample preparation with minimal hands-on effort. In contrast, workflows such as autoSP3 prepare 96 samples up to digestion within 3.5 hours, enabling ∼300 samples per day under manual supervision, while post-processing is not included ^7^. Similarly, a recently published plasma proteomics pipeline could process up to 40 × 96-well plates in under 300 minutes by coordinating liquid handling systems, centrifuges, and PCR-cyclers ^6^. However, this approach remains labour-intensive and highly specialized for plasma proteomics, limiting its flexibility and generalizability. When comparing workflows, it is crucial to note that our platform offers several additional steps beyond the primary processes of protein reduction, alkylation, and digestion. These include protein and peptide concentration determination with subsequent sample normalization, as well as peptide clean-up and desalting, which are essential for maintaining high-quality results. Our platform operates completely hands-free, including overnight and weekend runs, enabling the processing of hundreds of samples per week.

With further integration of 96- or 384-well pipetting heads and adapting processing to shorten incubation times, throughput could scale to thousands of samples per week, advancing both efficiency and reproducibility in proteomic sample preparation.

Bottom-up proteomic workflows are often used to address a variety of biological questions and can assess various information such as qualitative and quantitative protein expression ^4,50–52^, post-translational modifications ^53^, protein structural alterations ^54^ or identification of binding sites ^55,56^. All these different applications pose different requirements for the sample preparation, making the according workflows highly complex and versatile. This adds unique challenges to automation initiatives. A compelling example is the quantification of proteins using heavy-labelled reference peptides. Following protein identification, this approach requires spiking synthetic peptides into the digested sample at precisely defined amounts ^50^.

Because the reference-based quantification directly depends on the spiked-in volume, even minor inaccuracies during this step can lead to significant errors. Therefore, the automation system must support microliter pipetting precision and maintain consistency across large-scale high-throughput operations. Beyond peptide spike-ins, modern proteomics workflows often require additional modules tailored to specific experimental goals. This includes depletion of high-abundance proteins (e.g., in plasma or CSF) ^57^, proteome-wide enrichment of proteins ^58^, peptides ^59^ or modified peptides (PTMs) ^60,61^ and targeted protein and peptides capture strategies such as pull-downs or immunoprecipitations ^62^. Furthermore, recent advances such as limited proteolysis-MS ^55^ or thermal proteome profiling ^63^ demand workflows capable of accommodating diverse sample inputs and proteome states. Our automated platform addresses these challenges through a modular, block wise architecture, where processing blocks can be rearranged or extended to suit specific applications. Each module operates independently, allowing for flexible configuration without compromising throughput or reproducibility. For instance, enrichment steps can be inserted pre- or post-digestion, depletion modules can precede lysis, and heavy-labelled peptide spiking can be integrated seamlessly after clean-up, all on the same instrument. This design also future-proofs the system, allowing rapid adaptation as new proteomic techniques emerge. As the field continues to evolve, driven by improvements in enrichment techniques or miniaturized sample formats, having a platform that can scale, reconfigure, and expand is crucial for keeping pace.

Taken together, our flexible end-to-end automated sample preparation workflow enables reproducible large-scale applications in human medical research, supporting both current and next-generation applications in bottom-up proteomics.

## Material & Methods

All materials used for automated sample preparation are listed in Supplementary Data 1. The remaining materials are described in Supplementary Data 2. Frozen cell pellets were procured from Ipracell and CLS Service, live cells for cell culture were obtained from Addex Bio (Supplementary Data 3). Patient-derived PBMCs samples were purchased from Ruwag. Compounds employed for cell treatment were sourced from a combination of commercial suppliers, including Cambridge Bioscience and MedChemtronica AB and through custom synthesis by Sygnature Discovery (Supplementary Data 4). Heavy-labelled reference peptides (Supplementary Data 5) were synthesized by SciTide.

### Cell culture

A549 cells were cultured in Dulbecco’s Modified Eagle’s Medium (DMEM), supplemented with 10% heat-inactivated fetal bovine serum (HI FBS) and 1% Penicillin-Streptomycin solution. K562 cells were cultured in RPMI-1640 medium, supplemented with 10% fetal bovine serum (FBS) and 1% Penicillin-Streptomycin solution.

To determine the optimal cell density for compound treatment, A549 and K562 cells were seeded in flat bottom or U-bottom 96-well plates, respectively, at the following cell densities: 500,000, 225,000, 150,000, 100,000 and 75,000 cells/well, using eight replicate wells/condition and in a final volume of 100 µL/well. Five hours after seeding the cells were washed twice in phosphate-buffered saline (PBS, 1X) and the cell pellets were frozen at -80°C.

### Cell treatment with molecular degraders

For compound treatment, eight molecular glues and two bifunctional degraders (10 compounds listed in Supplementary Data 2) were diluted in dimethyl sulfoxide (DMSO) and added to flat bottom and U-bottom 96-well plates using an Echo 550 (Labcyte) liquid handler. The A549 and K562 cells were then seeded at 225,000 cells/well in flat bottom or U-bottom 96-well plates, respectively, containing pre-dispensed compounds, in a final volume of 100 µL/well. Molecular glues and bifunctional degraders were tested at the final concentration reported in Supplementary Data 4. The treatment was performed following a randomized plate map including five replicates/compound and 5 vehicle (DMSO) replicates/plate. The final concentration of DMSO was 0.1% in each well. After a five-hour treatment, the cells were washed twice in PBS, leading to a residual PBS volume in each well lower than 20 µL, and the cell pellets were frozen at -80°C prior to processing for mass spectrometry.

### Pre-processing – Cell lysis with ultrasonication

Frozen cell pellets (Supplementary Data 3) were resuspended in Biognosys lysis buffer and transferred to a 96 AFA-Tube TPX (Covaris) plate. The samples were lysed using sonication in a focused ultrasonicator LE220Rsc (Covaris) with processing duration 300 s per column, peak incident power of 450 W, duty factor of 35%, 200 cycles per burst, and Y-axis dithering of +/- 3 mm at 20 mm/s. Sonicated samples were incubated for 5 min at 90°C with 800 rpm followed by a centrifugation step for 2 min at 1000xg, 4°C. Protein concentration of the samples was determined using a bicinchoninic acid assay (BCA) assay kit (Thermo Fisher Scientific), following the manufacturer’s instructions.

### Automated sample preparation workflow

Automated sample preparation was performed using a custom-designed workcell centered around a Microlab STAR Plus (Hamilton) liquid handling system. The system incorporates a range of labware, accessories, and integrated third-party devices, all controlled by a suite of dedicated software packages. A detailed schematic of the system and an in-depth deck layout are provided in Supplementary Fig. 1 and 2, respectively. A complete list of hardware and associated software is available in Supplementary Data 1. Used labware to process the samples are listed in Supplementary Data 1.

#### Preparation

The main workflow encompasses five key processing blocks: (1) protein concentration determination, (2) protein aggregation capture (PAC) protocol, (3) peptide cleanup, (4) peptide concentration determination, and (5) LC-MS preparation. The workflow can be started at any of these blocks, selecting the appropriate starting point based on sample type and processing requirements. A sample worklist containing all necessary sample information must be provided to start the workflow. The system accommodates 1 to 192 samples, processed in one or two 96-well plates concurrently. Prior to execution, dynamic scheduling optimizes hardware resource allocation and maximizes throughput based on the number of samples. Samples can be loaded in tubes or 96-well plates and are subsequently cross-referenced with barcodes, if available. Samples loaded in tubes are then transferred to 96-well plates.

#### Protein concentration determination

Protein concentration was determined using a BCA protein assay kit according to the manufacturer’s instruction. Prior to the assay, lysed samples were equilibrated to room temperature on a thermoshaker for 15 minutes. Samples were then diluted 1:5 with PBS. For samples with initial concentrations exceeding the upper limit of the standard curve, another 1:5 dilution of diluted samples was performed, and the assay was repeated. The final protein concentration was calculated by multiplying the measured concentration by the appropriate dilution factor.

#### Protein aggregation capture (PAC)

Unless otherwise indicated, 75 µg of protein were transferred and concentrations normalized using Biognosys lysis buffer. The PAC protocol was performed as described previously ^12^. In brief, the sample was mixed with carboxylate-modified Sera-Mag SpeedBead magnetic particles (Sigma-Aldrich) and protein aggregation was induced by the addition of methanol for a final concentration of 70% (w/w). The suspension was mixed for 22 min at room temperature and the magnetic beads were retained on a magnetic rack for 90 sec. The beads were washed twice with 95% acetonitrile solution and once with 70% ethanol. Proteins were reduced and alkylated using reduction/alkylation buffer (0.1 M ammonium bicarbonate (ABC), 20 mM tris(2-carboxyethyl)phosphine (TCEP), 80 mM chloroacetamide (CAA), 70% ethanol) for 60 min at 37 °C. The supernatant was removed using the magnetic rack and the samples were digested with digestion buffer (50 mM tris(hydroxymethyl)aminomethane (Tris), trypsin (1:50 enzyme to substrate ratio), Lys-C (1:200 enzyme to substrate ratio)) for 5 hours at 37 °C. The beads were removed using a magnet, the supernatant was transferred to a new plate, and the samples were acidified with digestion stop buffer (20% trifluoroacetic acid (TFA)).

#### Peptide clean-up

Peptide clean-up and desalting were performed using positive pressure and solid-phase extraction (SPE) using hydrophilic-lipophilic balanced (HLB) 96-well plates (Waters). Prior to sample loading, the HLB plates were conditioned by sequentially adding 100% acetonitrile (ACN) and washing buffer (1% ACN, 0.1% TFA in water). Peptide samples were loaded onto the conditioned HLB plate washed twice with washing buffer (1% ACN, 0.1% TFA in water)., Peptides were eluted twice from the HLB plate using elution buffer (30% ACN, 0.1% TFA in water) and dried to completion using blowdown evaporation with heated nitrogen. Dried peptides were subsequently resolubilized in resuspension buffer (1% ACN, 0.1% FA in water with iRT peptides).

#### Peptide concentration determination

Peptide concentration was determined using a micro BCA (mBCA) protein assay kit (Thermo Fisher Scientific) according to the manufacturer’s instruction. For samples with initial concentrations exceeding the upper limit of the standard curve, a 1:5 dilution of the samples was performed, and the assay was repeated. The final peptide concentration was calculated by multiplying the measured concentration by the appropriate dilution factor.

#### Finalization

For final LC-MS preparation, resuspended samples were transferred to an LC-compatible output plate and all concentrations normalized to a user-defined value with resuspension buffer (0.1% formic acid spiked with iRT kit). The plate was sealed using a plate sealer. All generated data, including plate reader output files, trace and log files from hardware components, sample overview files were transferred to user-specified data storage location.

### Post-processing – heavy-labelled reference peptide spike-in

Following automated sample preparation workflow on the liquid handling system, reference-based quantification was performed. An equimolar master mix of twenty-one heavy-labelled peptides (Supplementary Data 5, SciTide) was prepared according to manufacturer’s instructions. Samples were normalized to the lowest peptide concentration of all samples using resuspension buffer (0.1% formic acid spiked with iRT kit) and then spiked with the heavy-labelled peptide master mix at a concentration of 8 fmol/µg endogenous peptides.

### Semi-automated sample preparation workflow

Frozen HeLa cell pellets were lysed with ultrasonication protocol, as described above. The resulting lysates were processed using a semi-automated PAC protocol implemented on a KingFisher Flex device (Thermo Fisher Scientific), following the general procedure outlined above ^12^. Digestion was performed overnight (18 h) on a thermomixer at 800 rpm, 37°C followed by separation of peptides and magnetic beads. Finally, the digestion was quenched by acidification with 20% TFA, ensuring a pH below 2. The resulting samples were processed with manual clean-up protocol (see below).

### Manual sample preparation

Frozen HeLa cell pellets were resuspended in lysis buffer (8 M urea) and then vortexed for 30 sec. Protein concentration of the samples was determined using BCA assay kit, according to the manufacturer’s instructions. Samples were reduced and alkylated (final concentration: 10 mM TCEP, 40 mM CAA) for 60 min at, 800 rpm, 37°C). Digestion was performed by diluting the samples with digestion buffer (0.1 M ABC, trypsin at a 1:50 enzyme to substrate ratio) to reach < 2 M Urea concentration followed by an overnight incubation (18 h) at 800 rpm, 37°C. Digestion was stopped by acidification with TFA (20%). Samples were further subjected to manual peptide clean-up.

### Manual peptide clean-up

Acidified peptides were purified by SPE with an HLB 96-well plate (Waters) according to the manufacturer’s instructions. Subsequently, elution buffer was removed using a SpeedVac vacuum concentrator (Thermo Fisher Scientific). Dried samples were then resuspended in resuspension buffer (0.1% formic acid spiked with iRT kit). Peptide concentrations were assessed with mBCA Protein assay kit according to the manufacturer’s instructions.

### Targeted assay development

The list of target peptides for LC high-field asymmetric-waveform ion mobility spectrometry coupled to parallel reaction monitoring (LC-FAIMS-PRM) analysis was created from FAIMS-DIA LC-MS/MS duplicate measurements of A549 and K562 peptide samples (225’000 starting cell amount). DIA analysis data was searched using the parameters described below with identified peptides displayed in Supplementary Figures 19-25. The best performing three proteotypic peptides per protein (Supplementary Data 5) were selected using SpectroDive 12 (Biognosys) interface. Peptides were selected by balancing amino acid sequence penalties (amino acids, i.e. Cysteine, Methionine), MS2 intensity, peptide rank, retention time, and missed cleavages, elaborated by Pauletti *et al.* ^64^. Selected peptides were synthesized with a AAA stable isotope label variant and subsequently used for heavy-labelled reference peptide spike-in, as described above.

### Targeted assay characterization

Peptides from A549 and K562 cells treated with a vehicle were utilized to establish a seven-point calibration curve, along with a blank, in triplicate injections. Each injection on the column contained 1 µg of peptides from either A549 or K562 cells. The concentration of heavy-labelled reference peptides ranged from 4 fmol to 0.000256 fmol per injection, with five-fold dilution steps in between.

The lower limit of quantification (LLOQ) for PRM assays was determined based on the following criteria: each replicate within the calibration point must have a q-value below 0.05, precision variation must be under 20%, accuracy must be better than 20% among dilution points, and intensity must be three times higher than the blank sample. The best-performing single peptide (from three available) per protein was selected for further sample analysis (Supplementary Data 6). Control protein PGK1 was used for normalization of PRM data across samples.

### Mass spectrometric acquisition

For FAIMS-DIA LC-MS/MS measurements (Fig. 2), samples with a high protein input (>30 µg) were normalized to an injection amount corresponding to peptide yields above 3.5 µg. For samples with a lower protein input (<30 µg), the entire sample material was injected, with volume normalization applied to ensure consistency. The samples were injected onto an in-house packed reversed-phase column using an EASY-nLC 1200 nano-LC system, interfaced with an Orbitrap Exploris 480 mass spectrometer equipped with a Nanospray Flex ion source (all from Thermo Fisher Scientific), a column oven (Sonation GMBH), and a FAIMS Pro ion mobility device (Thermo Fisher Scientific) operated at standard resolution (3.7 L/min carrier gas flow). LC solvents were A: water with 0.1 % FA; B: 80 % acetonitrile, 0.1 % FA in water. The nonlinear LC gradient was 1 - 50% solvent B in 171 minutes followed by a column washing step in 90% B for 7 minutes, and a final equilibration step of 1% B for 0.5 column volumes with a flow rate set to a ramp between 450 to 271 nL/min (min 0: 450 nL/min, min 172: 271 nL/min, washing at 400 nL/min). The FAIMS-DIA method consisted of one full range MS1 scan and 34 DIA segments per applied compensation voltage as described previously in Bruderer et al. ^65,66^.

For the measurements of negative controls (Fig. 2), 10 µL of sample were injected on an ACQUITY UHPLC CSH C18 reversed phase column (300 µm inner diameter, 15 cm length, 1.7 µm beads) on an Acquity UHPLC M-Class capillary flow liquid chromatography system (both Waters) connected to an Orbitrap Exploris 480 mass spectrometer operated in positive mode equipped with an EASY-spray ion source and a FAIMS Pro ion mobility device (all Thermo Fisher Scientific). The nonlinear LC gradient was 1 - 50% solvent B in 30 minutes followed by a column washing step in 90% B for 3 minutes, and a final equilibration step of 1% B for 3 minutes at 50°C with a flow rate set to 4 µL/min. The FAIMS-DIA method consisted per applied compensation voltage of one full range MS1 scan and 40 DIA seg- ments as adopted by Bruderer et. al ^65,66^.

For dia-PASEF LC-MS/MS acquisitions (Fig. 3, 4, 5) with 800 ng of peptides per sample were injected on an Aurora-3 series Ultimate CSI 75 µ m x 250 mm C18 reversed phase column (IonOpticks) on an EASY-nLC 1200 nano-LC system (Thermo Fisher Scientific) connected to a timsTOF HT mass spectrometer equipped with a Captive Spray II ion source (both Bruker Daltonics). LC solvents were A: water with 0.1% FA; B: 80% acetonitrile, 0.1% FA in water. The nonlinear LC gradient was 1 – 45% solvent B in 18 minutes followed by a column washing step in 90% B for 2.5 minutes, and a final equilibration step of 1% B for 3 minutes at 60 °C with a flow rate set to a ramp between 600 to 400 nL/min (min 0: 600 nL/min, min 18: 400 nL/min, washing at 600 nL/min). The dia-PASEF method consisted of one full range MS1 scan from 100 – 1700 m/z with an applied ion mobility range from 0.85 –1.45 1/k0 with ramp and accumulation times set to 70 ms (100% duty cycle) and 12 PASEF ramps.

For LC-FAIMS-PRM measurements (Fig. 5), 1 µg of peptides per sample were injected on an Aurora-3 series frontier-type CSI 75 µm x 550 mm C18 reversed phase custom proprietary column (IonOpticks) on a Vanquish Neo nano-LC system connected to a Exploris 480 mass spectrometer equipped with a FAIMS Pro device operated at standard resolution (3.7 L/min carrier gas flow) and connected to a Nanospray Flex source (all Thermo Fisher Scientific). LC solvents were A: 1% acetonitrile in water with 0.1% FA; B: 20% water in acetonitrile with 0.1% FA. The LC gradient was 0 – 50% Solvent B in 55 min followed by 50 - 90% B in 10 s, 90 % B for 8 min, with a flow rate set to a ramp between 500 to 200 nL/min (total gradient length was 63 min). A run in DIA mode was performed before data acquisition for retention time calibration using Biognosys’ iRT concept ^66^. A run in single ion monitoring mode according to FAIMS Pro user guidebooks was performed before data acquisition to optimize each precursor FAIMS Pro Compensation voltage, covering a range between -20 V and -80 V. The data acquisition window per peptide was 6 minutes. Target list is available in Supplementary Data 6.

### Mass spectrometric data analysis

Spectronaut 18 (Biognosys AG) was used to process DIA data files for Figures 2–4, Spectronaut 19 was used to process DIA data files for Figure 5 ^65^. directDIA analysis was run in pipeline mode using modified default settings. Specifically, the false discovery rate was set at 1% at the peptide spectrum match, peptide and protein group level with a maximal number of two missed cleavages and cross-run normalization using global normalization on the median. Trypsin was selected as digestion enzyme, with N-terminal protein acetylation and methionine oxidation set as variable modifications, and cysteine carbamidomethylation as a fixed modification. For the performance comparison evaluation, additional variable modifications of lysine and arginine carbamylation were included. Protein groups were defined as determined by the ID picker algorithm ^67^, which is implemented in Spectronaut. FASTA files were generated from UniProt sequences (Homo Sapiens, 2024-07-01, 20’435 entries) database. For the differential analyses, the precursor filtering was set to “Identified in % of the Runs (Percentile)” with value zero, as imputation strategy “Global Imputing” was selected and for the post analysis “Use All MS-Level Quantities” was enabled ^68^.

Signal processing and data analysis of PRM data were carried out using SpectroDive 12 (Biognosys) software based on mProphet algorithms ^69^. A q-value filter of 0.01 was applied for processed samples.

### Statistical data analysis

All statistical analyses were performed with the statistical language R version R-4.4.2 ^70^. Unless stated otherwise, functions from the stats version 4.2.2 package or the tidyverse version 2.0.0 package were used ^71^. Protein intensities were log2 transformed for further analysis.

For candidate ranking of differential abundance analysis of compound-treated vs. vehicle-treated samples, proteins with log FC > 0.58 were filtered and subsequently ranked by q-value. Peptide mapping was performed in PyMOL version 2.4.1 ^72^.

## Acknowledgements

The authors gratefully acknowledge Hamilton Bonaduz AG for their contributions to system design, initial method implementation and programming, and ongoing application support. We would also like to thank Matthew Carr at Sygnature Discovery for his assistance with the Echo protocol design.

## Author information

These authors contributed equally: Sandra Schär, Luca Räss.

## Authors and Affiliations

**Biognosys AG, Wagistrasse 21, 8952 Schlieren, Switzerland**

Sandra Schär, Luca Räss, Liliana Malinovska, Simonas Savickas, Christopher Below, Marco Tognetti, Polina Shichkova, Jakob Vowinkel, Yuehan Feng, Roland Bruderer and Lukas Reiter.

**Sygnature Discovery, Pennyfoot St, Nottingham NG1 1GR, United Kingdom** Francesca Cavallo, Benoit Gourdet, Gonzalo Robles, Leo Iu and Roland Hjerpe.

### Contributions

L.RÄ., designed, implemented, programmed and optimized the automated workflow. L.RÄ. and S.SC carried out all proteomics experiments and initial data analysis; S.SC., L.M. and P.S. carried out all downstream data analysis and visualization; C.B. established LC-MS methods; F.C. performed compound selection and cell treatment in cell-based assays; L.I. and G.R. performed compound synthesis; S.SA. carried out the targeted proteomics assay development, measurement and initial data analysis; M.T. established the initial workflow and system design. M.T., J.V., Y.F., R.H., B.G, R.B. and L.RE. provided project supervision; S.SC., L.RÄ. and L.M. conceived and designed the project and wrote the paper with input from all authors.

All authors have read and approved the manuscript.

### Corresponding authors

Correspondence to Roland Hjerpe, Roland Bruderer and Lukas Reiter.

## References

1. Macarron, R. et al. Impact of high-throughput screening in biomedical research. Nat. Rev. Drug Discov. 10, 188–195 (2011).

2. Schiff, L. et al. Integrating deep learning and unbiased automated high-content screening to identify complex disease signatures in human fibroblasts. Nat. Commun. 13, 1590 (2022).

3. Wenk, D., Zuo, C., Kislinger, T. & Sepiashvili, L. Recent developments in mass-spectrometry-based targeted proteomics of clinical cancer biomarkers. Clin. Proteomics 21, 6 (2024).

4. Jiang, Y. et al. Comprehensive overview of bottom-up proteomics using mass spectrometry. ACS Meas. Sci. Au 4, 338–417 (2024).

5. Manes, N. P. & Nita-Lazar, A. Application of targeted mass spectrometry in bottom-up proteomics for systems biology research. J. Proteomics 189, 75–90 (2018).

6. Albrecht, V., Müller-Reif, J. B., Brennsteiner, V. & Mann, M. A simplified perchloric acid workflow with neutralization (PCA-N) for democratizing deep plasma proteomics at population scale. bioRxiv (2025) doi:10.1101/2025.03.24.645089.

7. Müller, T. et al. Automated sample preparation with SP3 for low-input clinical proteomics. Mol. Syst. Biol. 16, e9111 (2020).

8. Dupree, E. J. et al. A critical review of bottom-up proteomics: The good, the bad, and the future of this field. Proteomes 8, 14 (2020).

9. Varnavides, G. et al. In search of a universal method: A comparative survey of bottom-up proteomics sample preparation methods. J. Proteome Res. 21, 2397–2411 (2022).

10. Duong, V.-A. & Lee, H. Bottom-up proteomics: Advancements in sample preparation. Int. J. Mol. Sci. 24, 5350 (2023).

11. Lee, J.-M., Han, J. J., Altwerger, G. & Kohn, E. C. Proteomics and biomarkers in clinical trials for drug development. J. Proteomics 74, 2632–2641 (2011).

12. Batth, T. S. et al. Protein aggregation capture on microparticles enables multipurpose proteomics sample preparation. Mol. Cell. Proteomics 18, 1027– 1035 (2019).

13. Medzihradszky, K. In-solution digestion of proteins for mass spectrometry. Methods Enzymol. 405, 50–65 (2005).

14. Kollipara, L. & Zahedi, R. P. Protein carbamylation: in vivo modification or in vitro artefact? Proteomics 13, 941–944 (2013).

15. Bhaskara, V. K., Mittal, B., Mysorekar, V. V., Amaresh, N. & Simal-Gandara, J. Resveratrol, cancer and cancer stem cells: A review on past to future. Curr. Res. Food Sci. 3, 284–295 (2020).

16. Harris, P. & Ralph, P. Human leukemic models of myelomonocytic development: a review of the HL-60 and U937 cell lines. J. Leukoc. Biol. 37, 407–422 (1985).

17. Torsvik, A. et al. U-251 revisited: genetic drift and phenotypic consequences of long-term cultures of glioblastoma cells. Cancer Med. 3, 812–824 (2014).

18. Thomas, P. & Smart, T. G. HEK293 cell line: a vehicle for the expression of recombinant proteins. J. Pharmacol. Toxicol. Methods 51, 187–200 (2005).

19. Alexovič, M. et al. Human peripheral blood mononuclear cells: A review of recent proteomic applications. Proteomics 22, e2200026 (2022).

20. Manning, G., Whyte, D. B., Martinez, R., Hunter, T. & Sudarsanam, S. The protein kinase complement of the human genome. Science 298, 1912–1934 (2002).

21. Liu, Y. et al. Expanding PROTACtable genome universe of E3 ligases. Nat. Commun. 14, 6509 (2023).

22. Zhang, M. et al. CDK inhibitors in cancer therapy, an overview of recent development. Am. J. Cancer Res. 11, 1913–1935 (2021).

23. Nishal, S., Jhawat, V., Gupta, S. & Phaugat, P. Utilization of kinase inhibitors as novel therapeutic drug targets: A review. Oncol. Res. 30, 221–230 (2022).

24. Drilon, A. et al. ROS1-dependent cancers - biology, diagnostics and therapeutics. Nat. Rev. Clin. Oncol. 18, 35–55 (2021).

25. Montor, W. R., Salas, A. R. O. S. E. & Melo, F. H. M. de. Receptor tyrosine kinases and downstream pathways as druggable targets for cancer treatment: the current arsenal of inhibitors. Mol. Cancer 17, 55 (2018).

26. Manea, C. A. et al. A review of NTRK fusions in cancer. Ann. Med. Surg. (Lond.) 79, 103893 (2022).

27. Kramer, L. T. & Zhang, X. Expanding the landscape of E3 ligases for targeted protein degradation. Curr. Res. Chem. Biol. 2, 100020 (2022).

28. Rodríguez-Gimeno, A. & Galdeano, C. Drug discovery approaches to target E3 ligases. Chembiochem 26, e202400656 (2025).

29. Quintás-Cardama, A. & Cortes, J. Molecular biology of bcr-abl1-positive chronic myeloid leukemia. Blood 113, 1619–1630 (2009).

30. Klein, E. et al. Properties of the K562 cell line, derived from a patient with chronic myeloid leukemia. Int. J. Cancer 18, 421–431 (1976).

31. Frere, G. A., de Araujo, E. D. & Gunning, P. Emerging mechanisms of targeted protein degradation by molecular glues. Methods Cell Biol. 169, 1–26 (2022).

32. Scholes, N. S., et al. Inhibitor-induced supercharging of kinase turnover via endogenous proteolytic circuits. bioRxiv (2024) doi:10.1101/2024.07.10.602881.

33. Jones, L. H. Small-molecule kinase downregulators. Cell Chem. Biol. 25, 30–35 (2018).

34. Tan, X., Huang, Z., Pei, H., Jia, Z. & Zheng, J. Molecular glue-mediated targeted protein degradation: A novel strategy in small-molecule drug development. iScience 27, 110712 (2024).

35. Békés, M., Langley, D. R. & Crews, C. M. PROTAC targeted protein degraders: the past is prologue. Nat. Rev. Drug Discov. 21, 181–200 (2022).

36. Feng, L., Zhang, H. & Liu, T. Multifaceted roles of IKZF1 gene, perspectives from bench to bedside. Front. Oncol. 14, 1383419 (2024).

37. Xia, R., Cheng, Y., Han, X., Wei, Y. & Wei, X. Ikaros proteins in tumor: Current perspectives and new developments. Front. Mol. Biosci. 8, 788440 (2021).

38. Cai, Y. & Wu, Y. The role of GSPT2 in tumor cell cycle regulation: Mechanisms and clinical significance. J. Cancer Ther. (2025) doi:10.4236/jct.2025.161002.

39. Zhang, D., Lin, P. & Lin, J. Molecular glues targeting GSPT1 in cancers: A potent therapy. Bioorg. Chem. 143, 107000 (2024).

40. Xiao, Y. & Dong, J. Coming of age: Targeting cyclin K in cancers. Cells 12, 2044 (2023).

41. Xu, Y., Nijhuis, A. & Keun, H. C. RNA-binding motif protein 39 (RBM39): An emerging cancer target. Br. J. Pharmacol. 179, 2795–2812 (2022).

42. Tolosa, E., Vila, M., Klein, C. & Rascol, O. LRRK2 in Parkinson disease: challenges of clinical trials. Nat. Rev. Neurol. 16, 97–107 (2020).

43. Ackerman, L. et al. IRAK4 degrader in hidradenitis suppurativa and atopic dermatitis: a phase 1 trial. Nat. Med. 29, 3127–3136 (2023).

44. Ye, X. et al. Combination of automated sample preparation and micro-flow LC-MS for high-throughput plasma proteomics. Clin. Proteomics 20, 3 (2023).

45. Burns, A. P. et al. A universal and high-throughput proteomics sample preparation platform. Anal. Chem. 93, 8423–8431 (2021).

46. Messner, C. B. et al. Ultra-high-throughput clinical proteomics reveals classifiers of COVID-19 infection. Cell Syst. 11, 11–24.e4 (2020).

47. Reder, A. et al. MassSpecPreppy-An end-to-end solution for automated protein concentration determination and flexible sample digestion for proteomics applications. Proteomics 24, e2300294 (2024).

48. Meier, F., Geyer, P. E., Virreira Winter, S., Cox, J. & Mann, M. BoxCar acquisition method enables single-shot proteomics at a depth of 10,000 proteins in 100 minutes. Nat. Methods 15, 440–448 (2018).

49. Guzman, U. H. et al. Ultra-fast label-free quantification and comprehensive proteome coverage with narrow-window data-independent acquisition. Nat. Biotechnol. 42, 1855–1866 (2024).

50. Lange, V., Picotti, P., Domon, B. & Aebersold, R. Selected reaction monitoring for quantitative proteomics: a tutorial. Mol. Syst. Biol. 4, 222 (2008).

51. Bian, Y. et al. Robust, reproducible and quantitative analysis of thousands of proteomes by micro-flow LC-MS/MS. Nat. Commun. 11, 157 (2020).

52. Bekker-Jensen, D. B. et al. An optimized shotgun strategy for the rapid generation of comprehensive human proteomes. Cell Syst. 4, 587–599.e4 (2017).

53. Hermann, J., Schurgers, L. & Jankowski, V. Identification and characterization of post-translational modifications: Clinical implications. Mol. Aspects Med. 86, 101066 (2022).

54. Malinovska, L. et al. Proteome-wide structural changes measured with limited proteolysis-mass spectrometry: an advanced protocol for high-throughput applications. Nat. Protoc. 18, 659–682 (2023).

55. Schopper, S. et al. Measuring protein structural changes on a proteome-wide scale using limited proteolysis-coupled mass spectrometry. Nat. Protoc. 12, 2391–2410 (2017).

56. Bludau, I. et al. Systematic detection of functional proteoform groups from bottom-up proteomic datasets. Nat. Commun. 12, 3810 (2021).

57. Tognetti, M. et al. Biomarker candidates for tumors identified from deep-profiled plasma stem predominantly from the low abundant area. J. Proteome Res. 21, 1718–1735 (2022).

58. Wu, C. C., et al. Mag-Net: Rapid enrichment of membrane-bound particles enables high coverage quantitative analysis of the plasma proteome. bioRxivorg (2024) doi:10.1101/2023.06.10.544439.

59. Poetz, O., Hoeppe, S., Templin, M. F., Stoll, D. & Joos, T. O. Proteome wide screening using peptide affinity capture. Proteomics 9, 1518–1523 (2009).

60. Batalha, I. L., Lowe, C. R. & Roque, A. C. A. Platforms for enrichment of phosphorylated proteins and peptides in proteomics. Trends Biotechnol. 30, 100–110 (2012).

61. Carlson, S. M., Moore, K. E., Green, E. M., Martín, G. M. & Gozani, O. Proteome-wide enrichment of proteins modified by lysine methylation. Nat. Protoc. 9, 37–50 (2014).

62. Anderson, N. L. et al. Mass spectrometric quantitation of peptides and proteins using Stable Isotope Standards and Capture by Anti-Peptide Antibodies (SISCAPA). J. Proteome Res. 3, 235–244 (2004).

63. Kurzawa, N. et al. Deep thermal profiling for detection of functional proteoform groups. Nat. Chem. Biol. 19, 962–971 (2023).

64. Pauletti, B. A. et al. Typic: A practical and robust tool to rank proteotypic peptides for targeted proteomics. J. Proteome Res. 22, 539–545 (2023).

65. Bruderer, R. et al. Extending the limits of quantitative proteome profiling with data-independent acquisition and application to acetaminophen-treated three-dimensional liver microtissues. Mol. Cell. Proteomics 14, 1400–1410 (2015).

66. Escher, C. et al. Using iRT, a normalized retention time for more targeted measurement of peptides. Proteomics 12, 1111–1121 (2012).

67. Ma, Z.-Q. et al. IDPicker 2.0: Improved protein assembly with high discrimination peptide identification filtering. J. Proteome Res. 8, 3872–3881 (2009).

68. Huang, T. et al. Combining precursor and fragment information for improved detection of differential abundance in data independent acquisition. Mol. Cell. Proteomics 19, 421–430 (2020).

69. Reiter, L. et al. mProphet: automated data processing and statistical validation for large-scale SRM experiments. Nat. Methods 8, 430–435 (2011).

70. Team, R. D. C. R: A language and environment for statistical computing. *(No Title)* (2010).

71. Wickham, H. et al. Welcome to the tidyverse. J. Open Source Softw. 4, 1686 (2019).

72. Schrodinger, L. L. C. The PyMOL molecular graphics system. *Version* (2015).

73. Deutsch, E. W. et al. The ProteomeXchange consortium at 10 years: 2023 update. Nucleic Acids Res. 51, D1539–D1548 (2023).

