## Supplementary Material for "A flexible end-to-end automated sample preparation workflow enables reproducible large-scale bottom-up proteomics"

#### Supplementary Results

##### Automated sample preparation coupled to LC-MS can be utilized for target identification, quantification and selectivity profiling of molecular degraders

We used the information from the cell line characterization (Fig. 4) to identify two suitable cell lines. Based on the expression level of the known targets, we selected K562 and A549 as the most suitable cell lines for the treatment (Supplementary Fig. 11). Next, we determined the optimal cell density for treatment conditions, balancing protein and peptide identifications. We tested cell counts of 500,000, 225,000, 150,000, 100,000, and 75,000 per well, and found stable protein group and peptide identification with low coefficients of variation (CVs) for each protein group quantity calculated across quadruplicates, all below 6.8% (Supplementary Fig. 12 – 13). For a cell count of 225,000 cells per well, we observed an average peptide yield of 18 µg, which is sufficient for all measurements, thus we decided on this cell density for the compound treatment screen (Supplementary Fig. 14).

Pomalidomide, Cemsidomide and Iberdomide all target the IKZF1 and IKZF3 transcription factors, and clear degradation of these proteins can be observed for all three compounds. Additionally, we found that Pomalidomide, Cemsidomide, and Iberdomide significantly reduce ZFP91 levels. MRT-2359 can be observed to clearly downregulate the levels of GSPT1 and the paralogue GSPT2 in K562 and A549 cells. Two reported degraders, XL01126 and Compound-68 from patent WO2022198112 of LRRK2 were tested. Whilst XL01126 recruits the VHL ligase, Compound-68 is a Cereblon-based degrader, purported to be the clinical stage compound ARV-102 or a close analogue thereof. Consistent with LRRK2 expression levels in the two cell-lines, detection of compound-induced degradation was only detectable in A549 cells. This is evident for both XL01126 and Compound-68. To confirm and extend our results, we conducted a quantitative analysis of LRRK2 protein levels. This analysis confirmed that Compound-68 and XL01126 effectively degraded 70% of LRRK2 peptides. Cyclin K (CCNK) is implicated in regulation of RNA polymerase II and transcription [1]. R-CR8 was demonstrated to be a molecular glue degrader of CCNK, and following this, other CCNK degraders, such as SR-4835, have been reported [2,3]. From our results, it is apparent that both R-CR8 and SR-

4835 treatments lead to significant changes in a large number of proteins, suggesting a potential transcriptional remodeling of the protein landscape in addition to targets degraded directly by the compounds. A relatively acute exposure time (5 h) was utilized to reduce the impact of transcriptional changes impacting protein abundance readouts, however the results indicate significant upregulation of several transcriptional regulators in A549 cells, including p53, FOS, FOSB, HES1 and HIF1A, which may in turn up- or down-regulate expression of a multitude of proteins. Notably, p53 was observed to be the top upregulated protein in this cell-line, with a log2 ratio of 2.6 and 2.9 for R-CR8 and SR-4835 treatment, respectively. In K562 cells, none of the above-mentioned transcription factors are identified as significantly regulated by the compounds. K562 cells harbours mutated and transcriptionally inactive p53, and the lower number of significantly regulated proteins in this cell line compared to A549 cells may be attributed to the inactive status of this transcription factor R-CR8 regulates 338 proteins in A549 cells vs. 214 in K562 cells; SR-4835 regulates 348 proteins in A549 cells vs. 225 in K562 cells [4]. In addition to these findings, CCNK was consistently and distinctly identified as a target of both R-CR8 and SR-4835 in both cell lines. The target quantity fell below the limit of quantification (BLQ), indicating a degradation of over 95%. The analogues E7070 and E7829 both target RBM39 and RBM23 via ternary the DCAF15 ubiquitin-ligase, and these proteins can be clearly observed to be downregulated in both A549 and K562 cells.

### Supplementary Figures

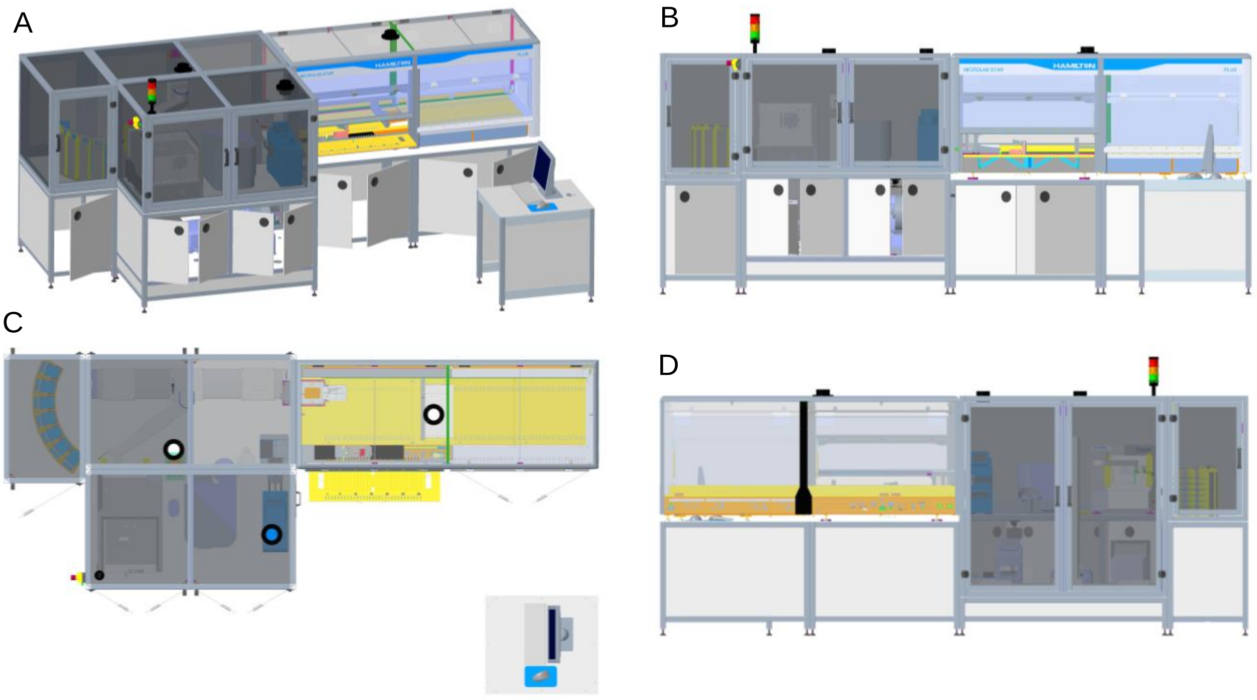


***Supplementary Figure 1 Graphical projections overview of the automated sample preparation system.*** *Detailed overview of the Hamilton STAR liquid handling robotic platform with integrated third-party devices. A) Front view from an elevated angle. B) Front view. C) Top-down view. Black rings depict air exhaust pipes. D) Back view.*

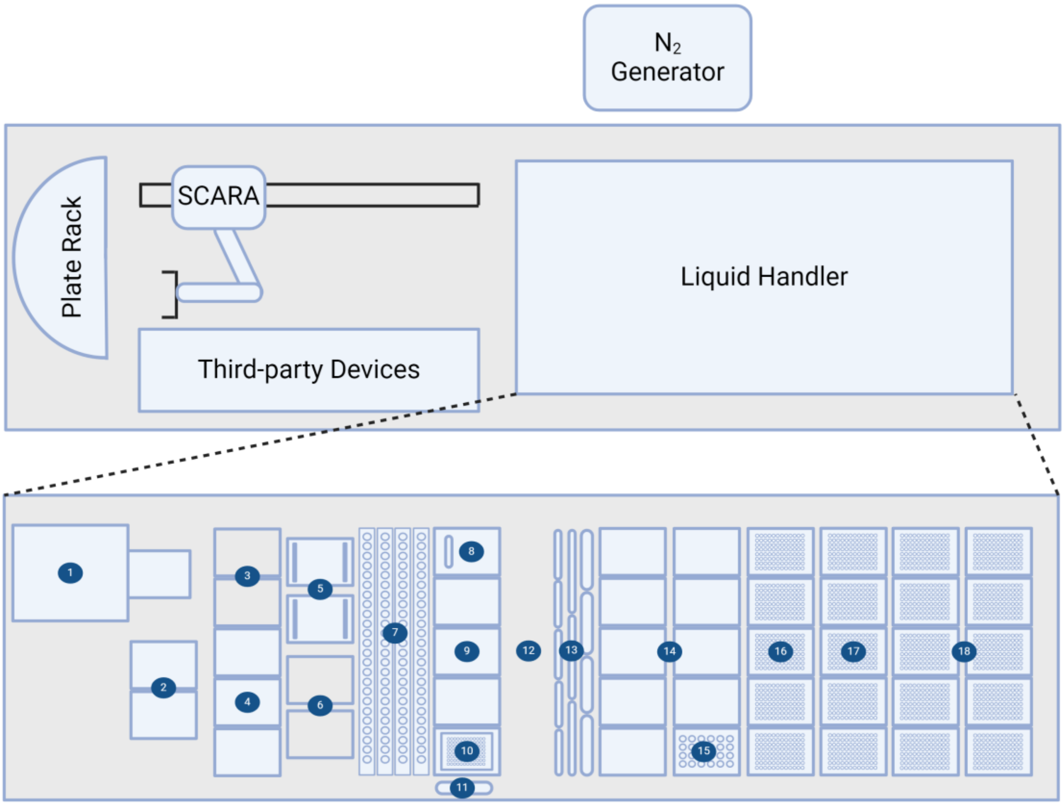


***Supplementary Figure 2 Layout of the automated sample preparation system and liquid handling system deck.*** *Top panel: Top-down view of the workcell layout illustrating the integration of the liquid handling system with various third-party devices and a plate storage rack enabled by a robotic SCARA arm. Third party devices*

*include an ultrasonicator, a plate sealer, a plate peeler, a microplate spectrophotometer and a nitrogen blowdown evaporator with its corresponding nitrogen generator. Lower panel: Deck layout of liquid handling system including (1) positive pressure extraction module, (2) 96-well handover positions accessible by SCARA arm as well as liquid handling system, (3) clean-up plate park position, (4) 96-well plate positions, (5) thermoshakers, (6) 1000 µL tip park positions, (7) tube carriers with 24 positions each, (8) refillable liquid reservoir, (9) 96-well plate positions, (10) 96-well magnet plate, (11) 1D barcode reader, (12) labware and liquid waste, (13) liquid reservoirs 60 mL, 120 mL and 200 mL, (14) temperature-regulated 96-well plate positions, (15) temperature-regulated tube carrier with 24 positions, (16) 1000 µL pipette tips, (17) stacked 50 µL pipette tips, (18) stacked 300 µL pipette tips. Created with Biorender.com.*

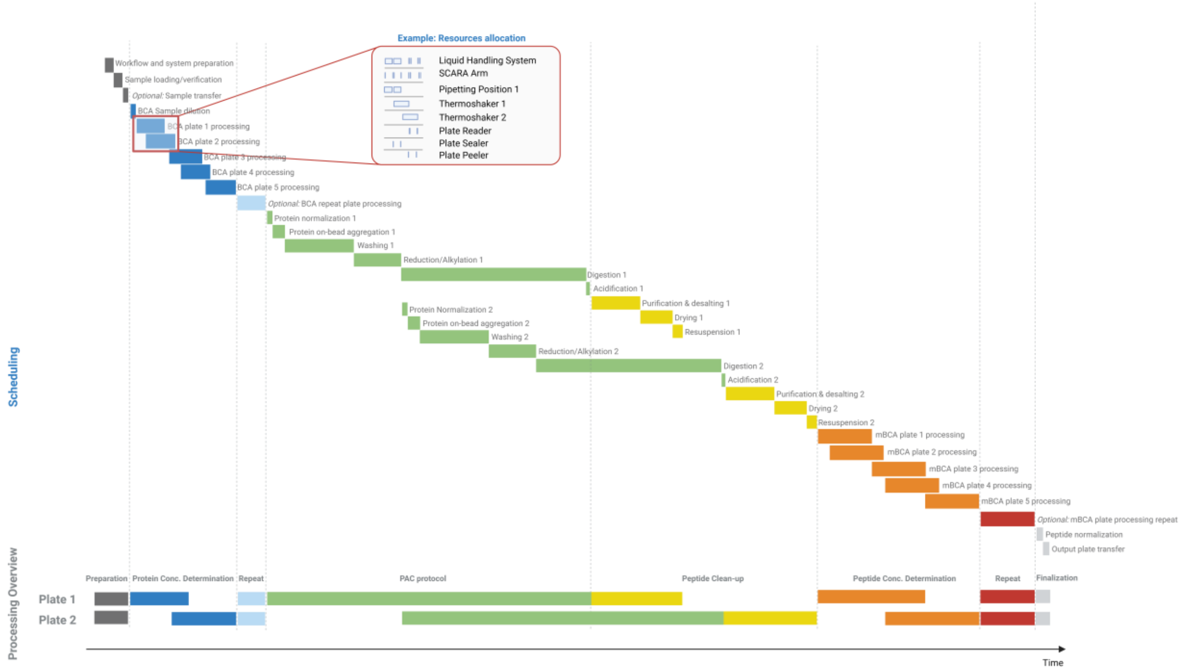


***Supplementary Figure 3 Workflow scheduling and resource allocation.** Figure illustrates the dynamic scheduling and resource allocation strategy for processing two 96-well plates. The top panel shows the planned sequence of processing steps, highlighting parallelization and resource allocation (inset). Box sizes are not representative. Dynamic scheduling optimizes the workflow in real-time based on resource availability, predicted processing times, and step dependencies. The bottom panel maps these steps to overarching processing blocks (dashed lines indicate block boundaries). Created with BioRender.com.*

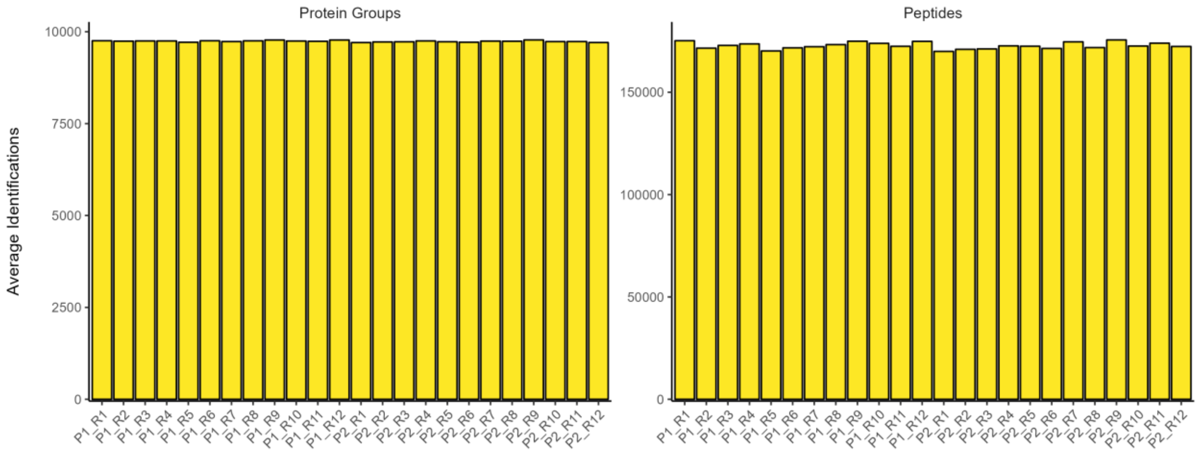


***Supplementary Figure 4 Assessment of reproducibility, sensitivity and precision of the automated sample preparation workflow.** Number of identified protein groups (left) and peptides (right) from the intra- and interplate reproducibility assessment across 24 replicates (R1 – R12) spanning two plates (P1 – P2).*

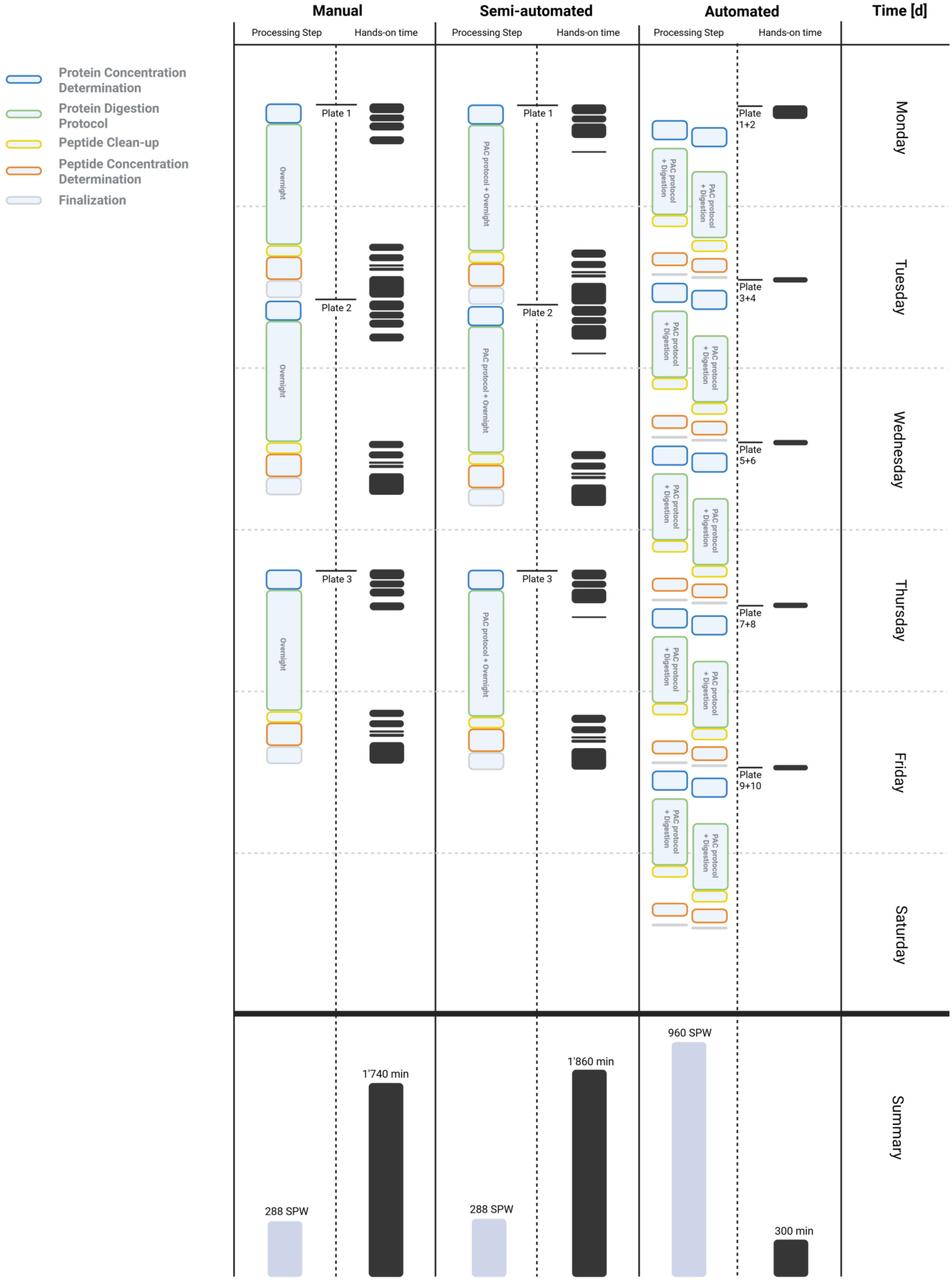

Protein Concentration Determination
Protein Digestion Protocol
Peptide Clean-up
Peptide Concentration Determination
Finalization
Manual
Semi-automated
Automated
Time [d]
Processing Step
Hands-on time
Plate 1
Plate 2
Plate 3
Overnight
PAC protocol + Overnight
PAC protocol + Digestion
Plate 1+2
Plate 3+4
Plate 5+6
Plate 7+8
Plate 9+10
Monday
Tuesday
Wednesday
Thursday
Friday
Saturday
Summary
288 SPW
1'740 min
288 SPW
1'860 min
960 SPW
300 min

***Supplementary Figure 5 Performance comparison between automated, semi-automated and manual sample processing workflow.*** *Processing time of 96-well plates for bottom-up proteomics sample preparation main steps, including protein concentration determination (blue), protein digestion protocol (green), peptide clean-up (yellow), peptide concentration determination (orange), and finalization (grey) for manual, semi-automated, and automated sample processing methods. User-dependent interventions for each processing step are depicted in black. The illustrated process spans from initial lysate input to the generation of concentration normalized peptide samples. The lower panel represents quantitative metrics of throughput, specifically illustrating the total number of samples processable per working week (SPW) and a consolidated representation of the cumulative user intervention time (black) across the three processing modalities. Created with BioRender.com.*

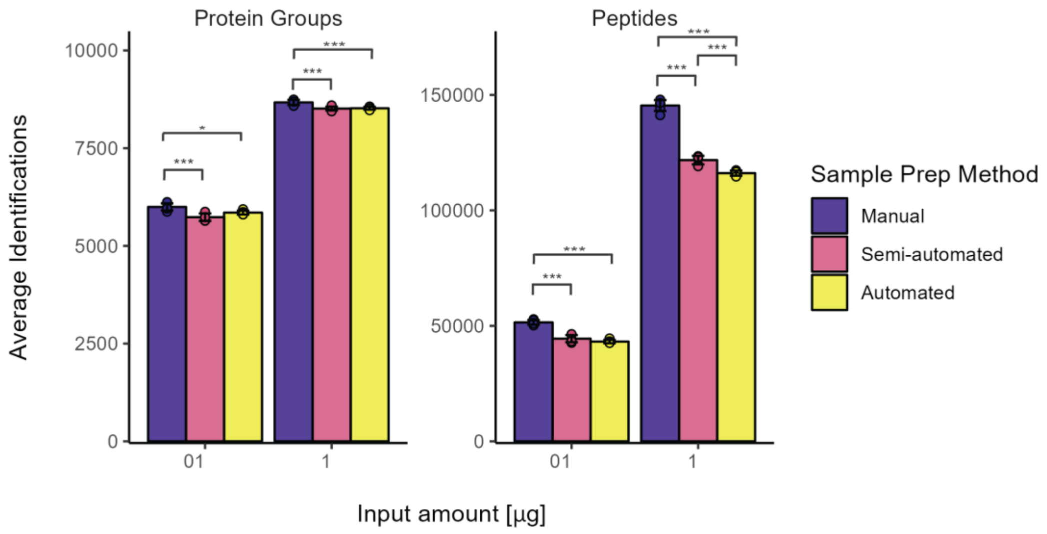


***Supplementary Figure 6 Performance comparison between automated, semi-automated and manual sample processing workflow.*** *Number of identified protein groups (left) and peptides (right) for protein input amounts of 0.1 and 1 µg, processed using the automated (yellow), the semi-automated (pink), and the manual (purple) workflow (n = 6, error bars: standard deviation). Statistical significance is determined by one-way ANOVA with Tukey's HSD: *p < 0.05, **p < 0.01, ***p < 0.001.*

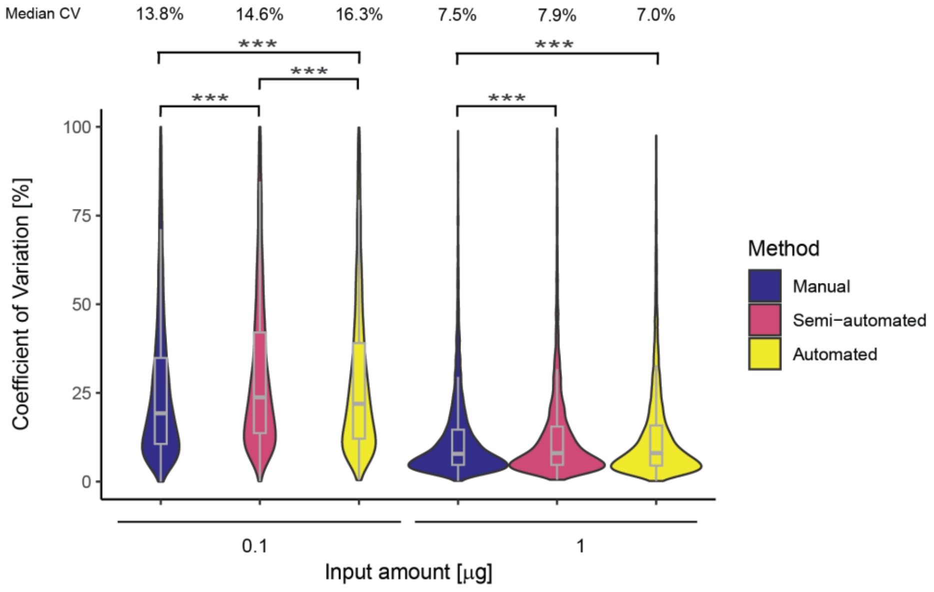


***Supplementary Figure 7 Performance comparison between automated, semi-automated and manual sample processing workflow.*** *Coefficient of variation (CV) of protein group quantities across the input amount of 0.1 and 1 µg protein, prepared using the automated (yellow), the semi-automated (pink), and the manual (purple) workflow (n = 6) depicted as violin plots. Embedded box plots display the median (thick lines), interquartile range (25th and 75th percentiles), and whiskers extending to ±1.58 times the interquartile range.*

*Significance groupings based upon one-way ANOVA with Tukey's HSD *p < 0.05, **p < 0.01, ***p < 0.001. Median CVs are shown above.*

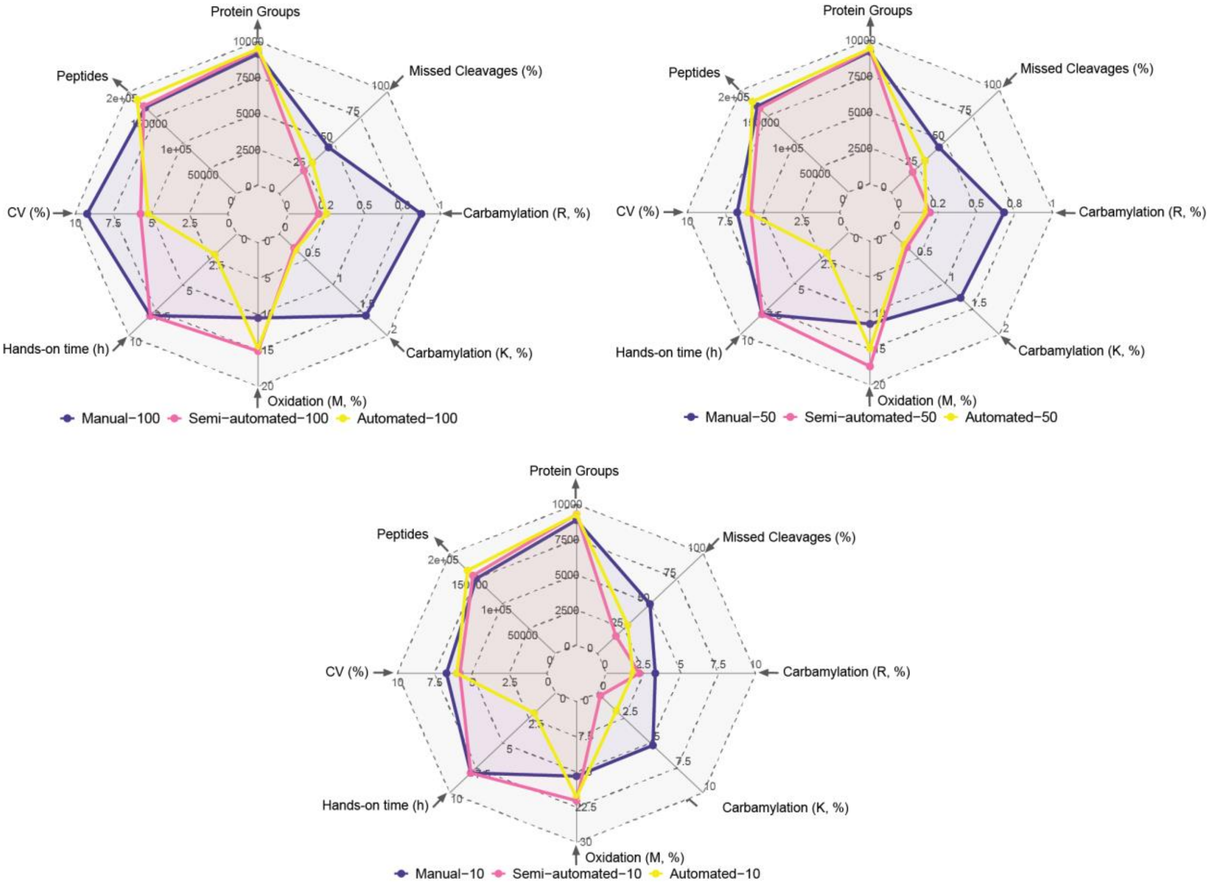


***Supplementary Figure 8 Performance comparison between automated, semi-automated and manual sample processing workflow.** Spider web plots visualizing number of protein groups and peptides, protein group CVs, hands-on time, relative proportion of post-translational modifications (e.g., oxidation, carbamylation of lysine (K) and arginine (R)) and missed cleavage ratio for 100 µg (top left), 50 µg (top right) and 10 µg (bottom) protein input amount prepared using the automated (yellow), the semi-automated (pink), and the manual (purple) workflow.*

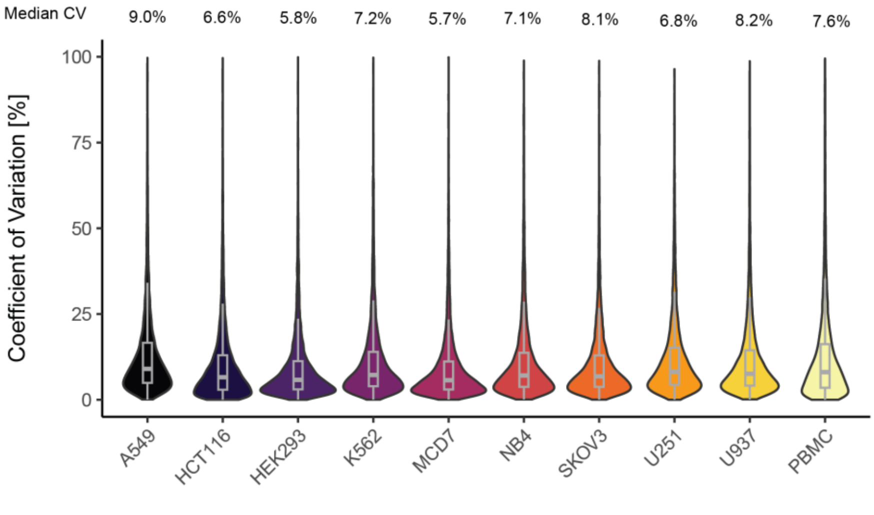


***Supplementary Figure 9 Characterization of multiple cell lines using the automated sample preparation system coupled to short-gradient LC-MS system**. Coefficient of variation (CV) of protein group quantities across cell line replicates depicted as violin plots. Embedded box plots display the median (thick lines), interquartile range (25th and 75th percentiles), and whiskers extending to ±1.58 times the interquartile range. Median CVs are shown above the violin plots.*

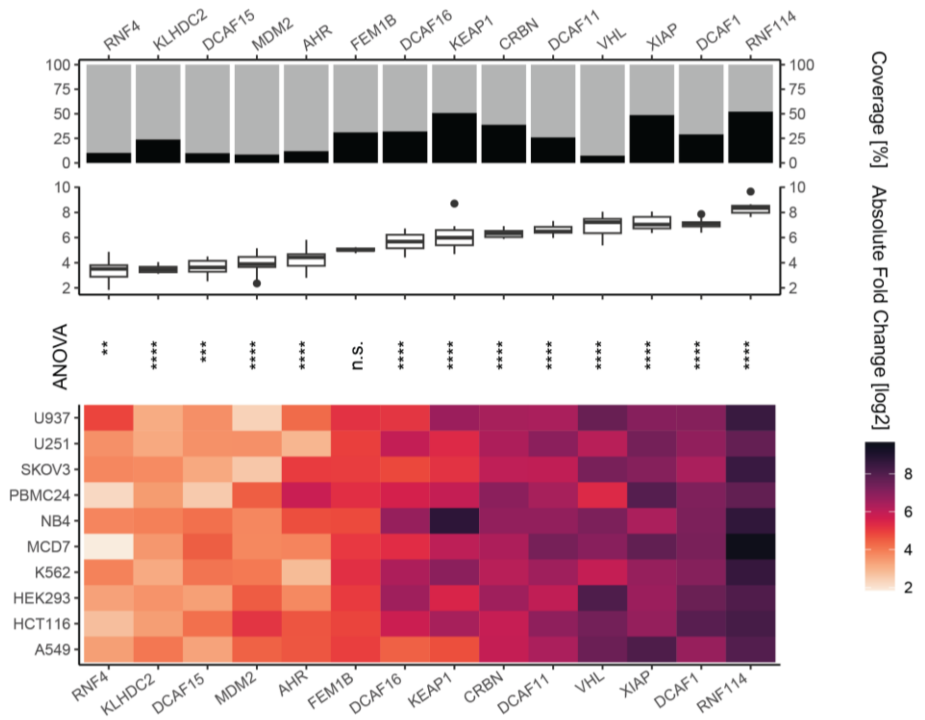


**Supplementary Figure 10 Characterization of multiple cell lines using the automated sample preparation workflow coupled to short-gradient LC-MS acquisition**. *Bar plots illustrate peptide coverage per selected E3 ligase across cell lines (top panel). Box plots show the median (thick lines), interquartile range (25th and 75th percentiles), and whiskers extending to ±1.58 times the interquartile range for the intensities of selected kinases across cell lines (middle panel). Heatmap displaying the mean protein intensities of selected E3 ligases across cell lines (lowest panel). One-way ANOVA tests for differences in mean protein intensities between cell lines. Significance labels derived from Tukey's HSD *p < 0.05, **p < 0.01, ***p < 0.001, ****p < 0.0001, n.s. indicates not significant.*

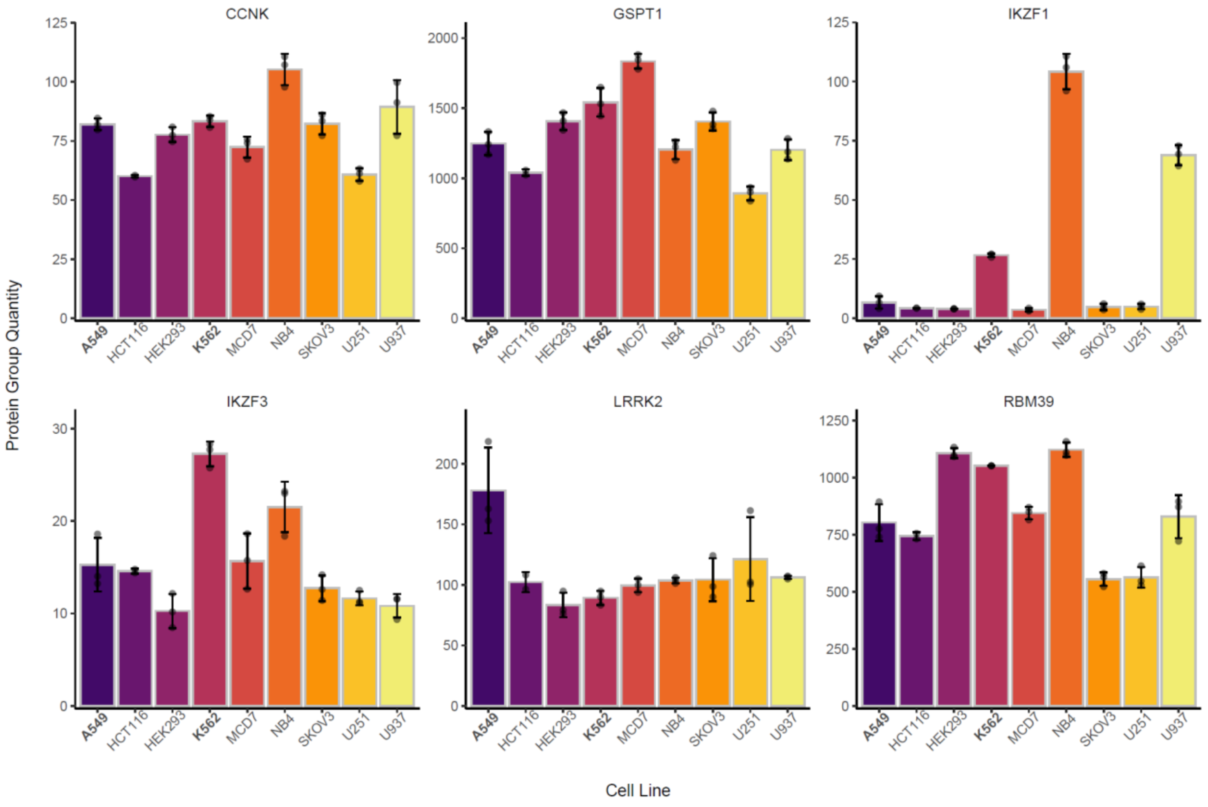


**Supplementary Figure 11 Automated sample preparation coupled to LC-MS can be utilized for target identification, quantification and selectivity profiling of molecular degraders.** *Protein group quantities of selected targets across ten cell lines (error bars: standard deviation).*

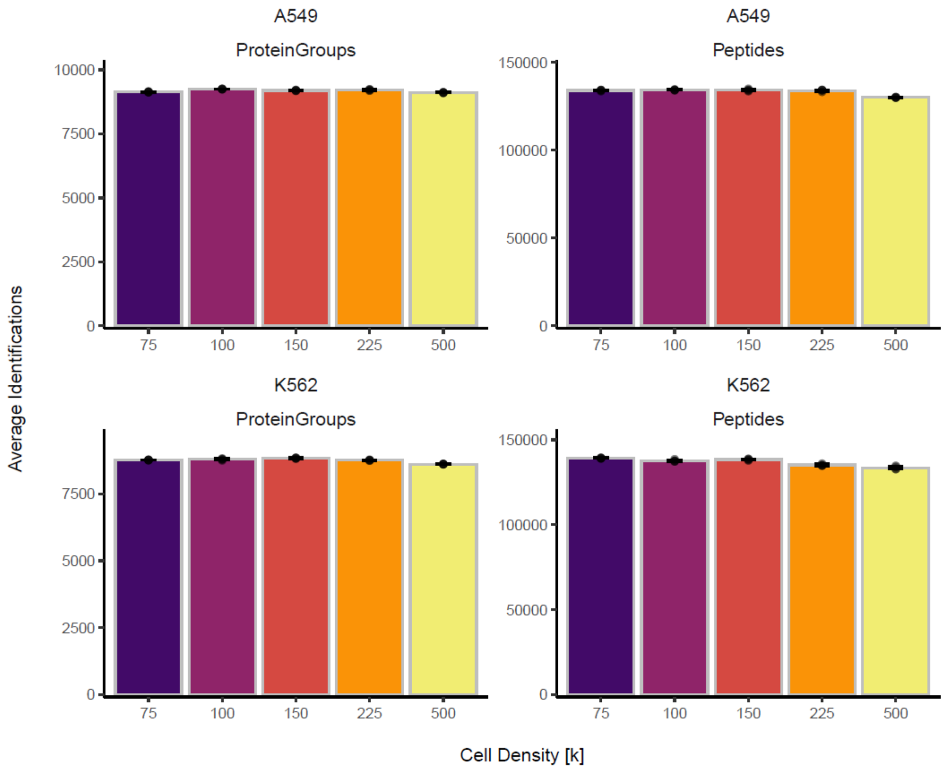


***Supplementary Figure 12 Automated sample preparation coupled to LC-MS can be utilized for target identification, quantification and selectivity profiling of molecular degraders****. Number of identified protein groups (left) and peptides (right) for cell densities of 75’000, 100’000, 150’000, 225’000 and 500’000 cells for two cell lines (A549 on top and K562 on the bottom), processed using the automated sample preparation workflow (n = 4, error bars: standard deviation).*

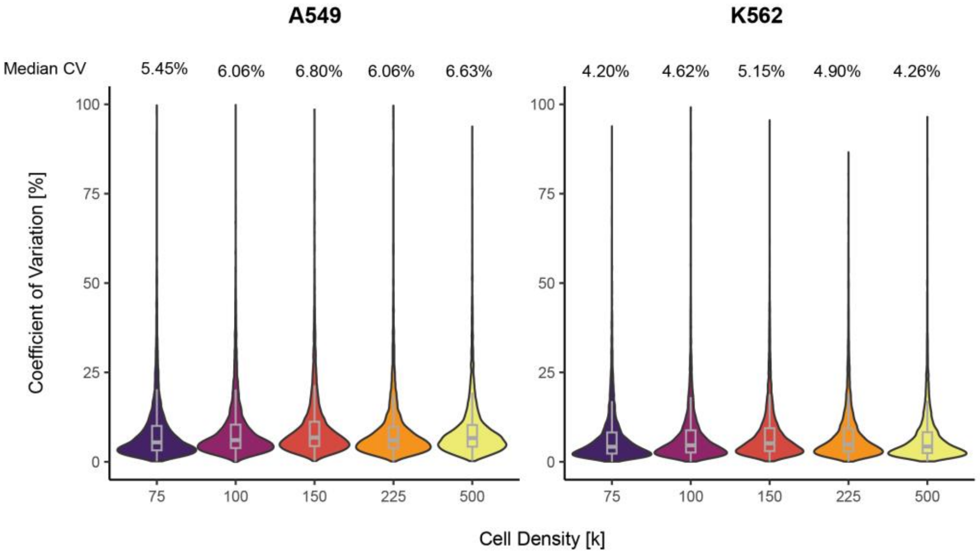


***Supplementary Figure 13 Automated sample preparation coupled to LC-MS can be utilized for target identification, quantification and selectivity profiling of molecular degraders.*** *Coefficient of variation (CV) of protein group quantities across cell densities of 75’000, 100’000, 150’000, 225’000 and 500’000 cells for two cell lines (A549 left and K562 right) (n = 4) depicted as violin plots. Embedded box plots display the median (thick lines), interquartile range (25th and 75th percentiles), and whiskers extending to ±1.58 times the interquartile range. Median CVs are shown above the violin plots.*

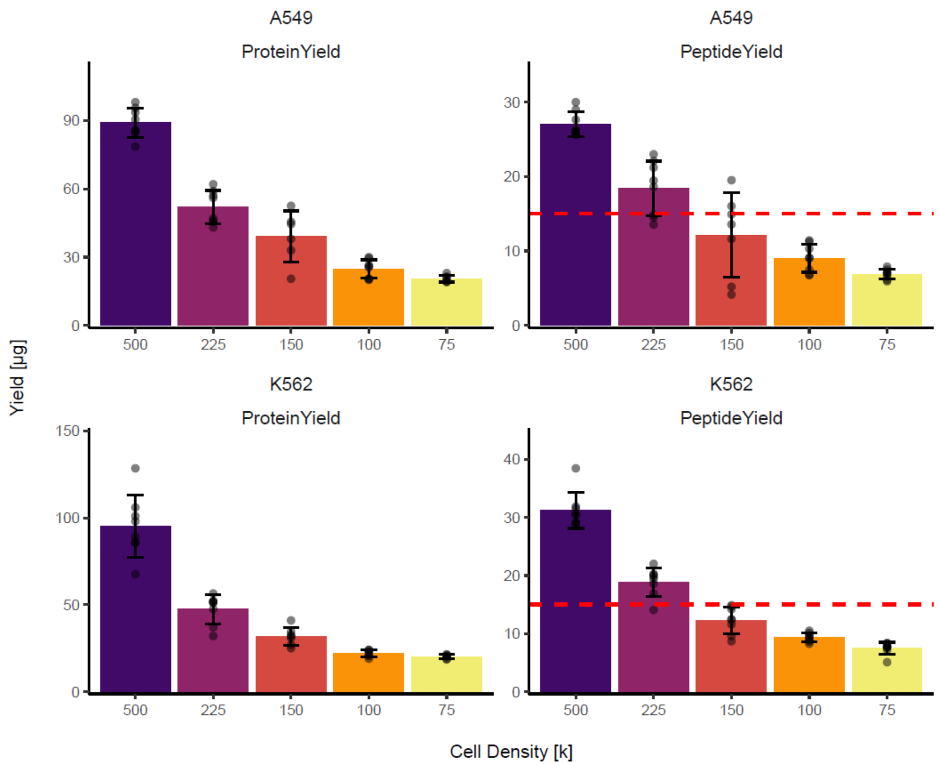


***Supplementary Figure 14 Automated sample preparation coupled to LC-MS can be utilized for target identification, quantification and selectivity profiling of molecular degraders.*** *Protein (left) and peptide (right) yields across cell densities of 75,000, 100,000, 150,000, 225,000, and 500,000 cells are shown for two cell lines: A549 (top barplots) and K562 (bottom barplots) (n = 8, error bars: standard deviation).*

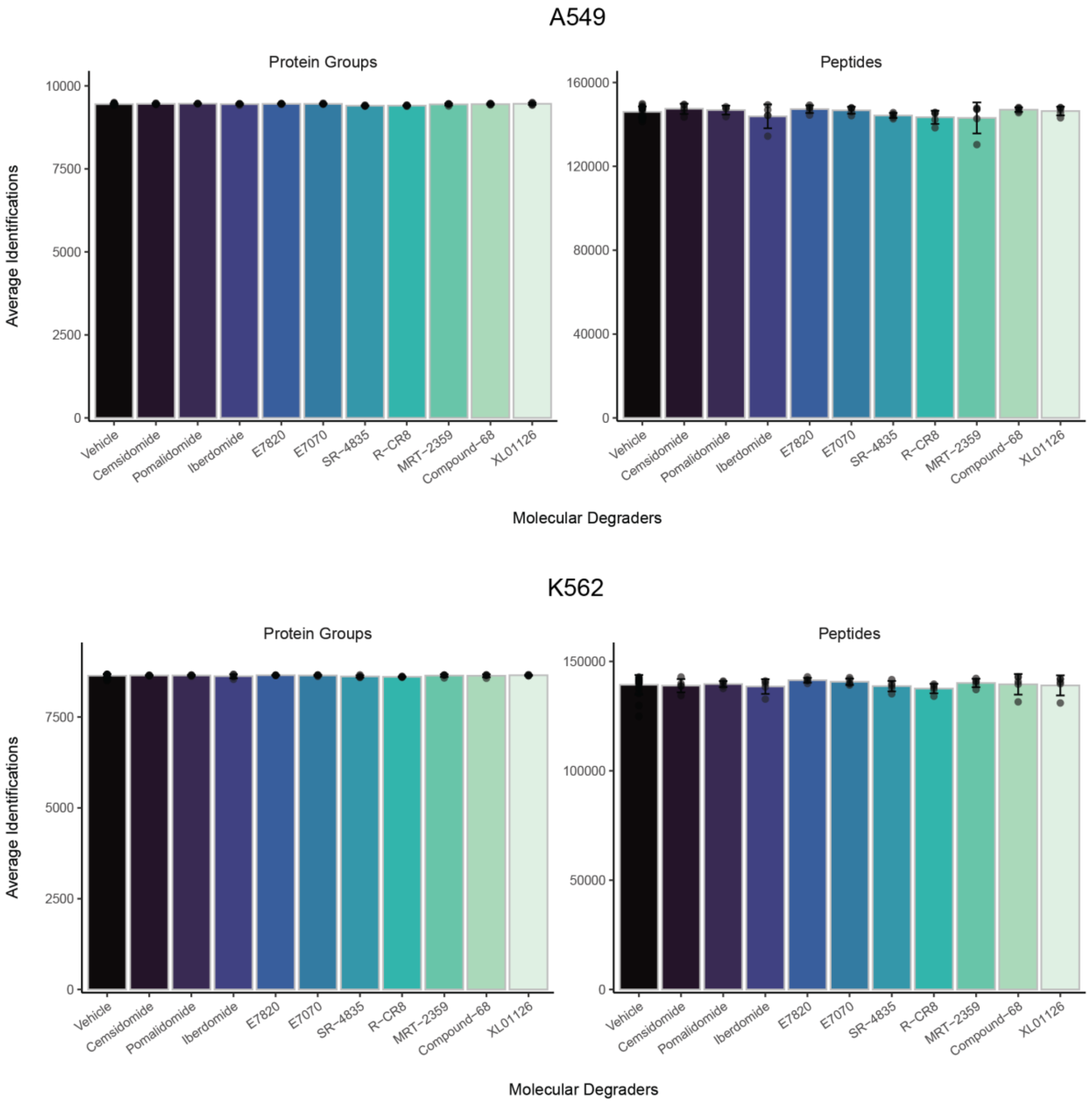


***Supplementary Figure 15 Automated sample preparation coupled to LC-MS can be utilized for target identification, quantification and selectivity profiling of molecular degraders.*** *Number of identified protein groups (left) and peptides (right) across 11 conditions, including one vehicle condition (DMSO) and ten conditions where cells were treated with distinct molecular degrader compounds (n = 5, error bars: standard deviation) on two cell lines, A549 (top) and K562 (bottom). Different conditions (vehicle and molecular degraders) are represented using a color gradient ranging from black to light green.*

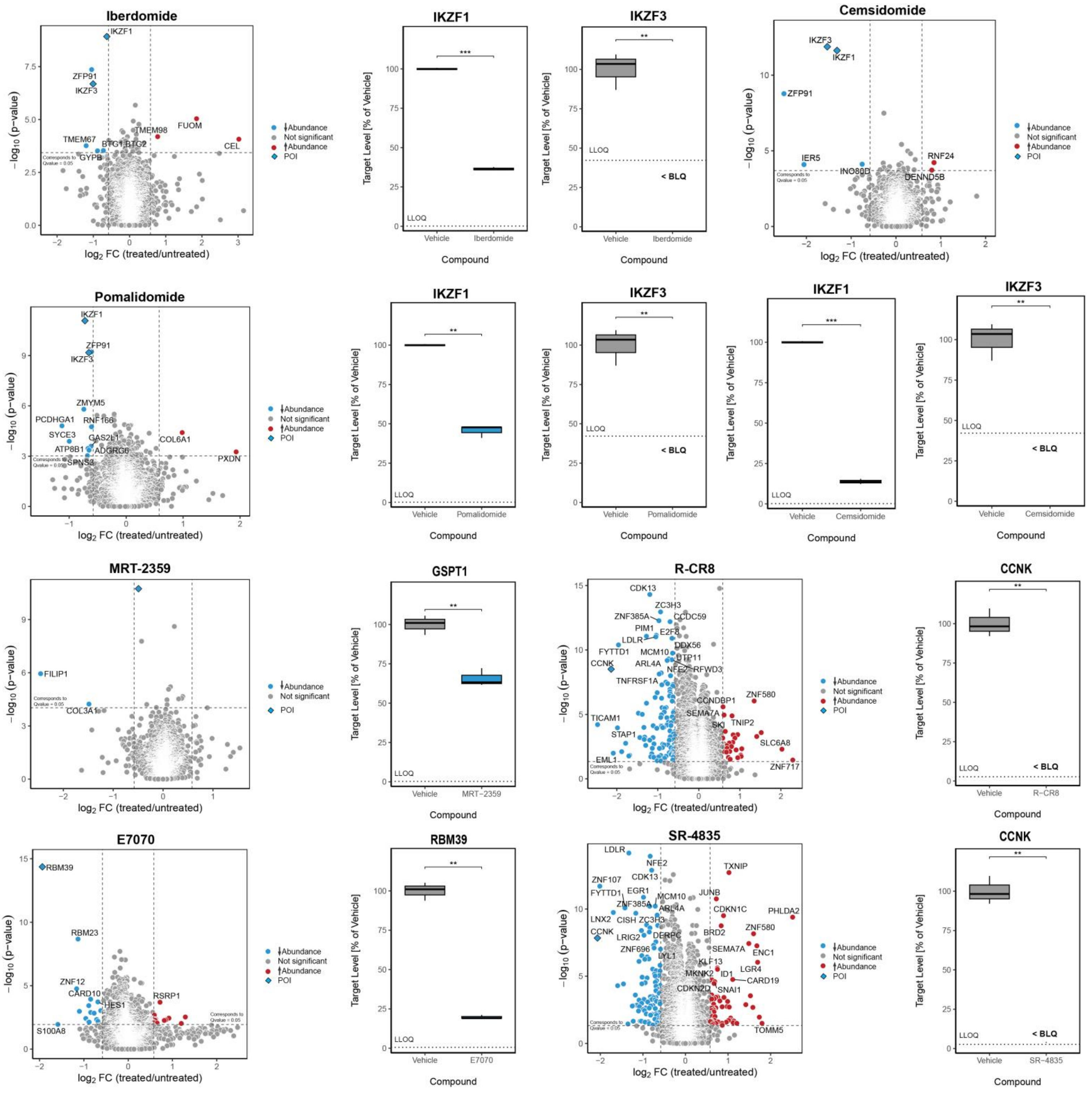


***Supplementary Figure 16 Automated sample preparation coupled to LC-MS can be utilized for target identification, quantification and selectivity profiling of molecular degraders.*** *Volcano plot illustrating the differentially abundant proteins identified through proteome profiling between K562 cell samples (n = 5) treated with the molecular degrader compounds (Supplementary Data 4, Supplementary Figure 17). Proteins are color-coded based on their statistical significance and fold change: red indicates increased abundance, blue indicates decreased abundance (|log2(FC)| > 0.58, q-value < 0.05) and grey indicates no significant change (|log2(FC)| ≤ 0.58, q-value > 0.05). Boxplot showing the absolute amount of the POIs in K562 cell samples (n = 3) treated with the molecular degrader compounds, compared to DMSO-treated samples (vehicle). T-test based p-values are indicated in the plots: * = p < 0.05, ** = p < 0.01, *** = p < 0.001, **** = p < 0.0001, n.s. = not significant. Lower limit of quantification (LLOQ) is indicated as dotted line. Targets falling below the lower limit of quantification (LLOQ) are marked as below limit of quantification (<BLQ).*

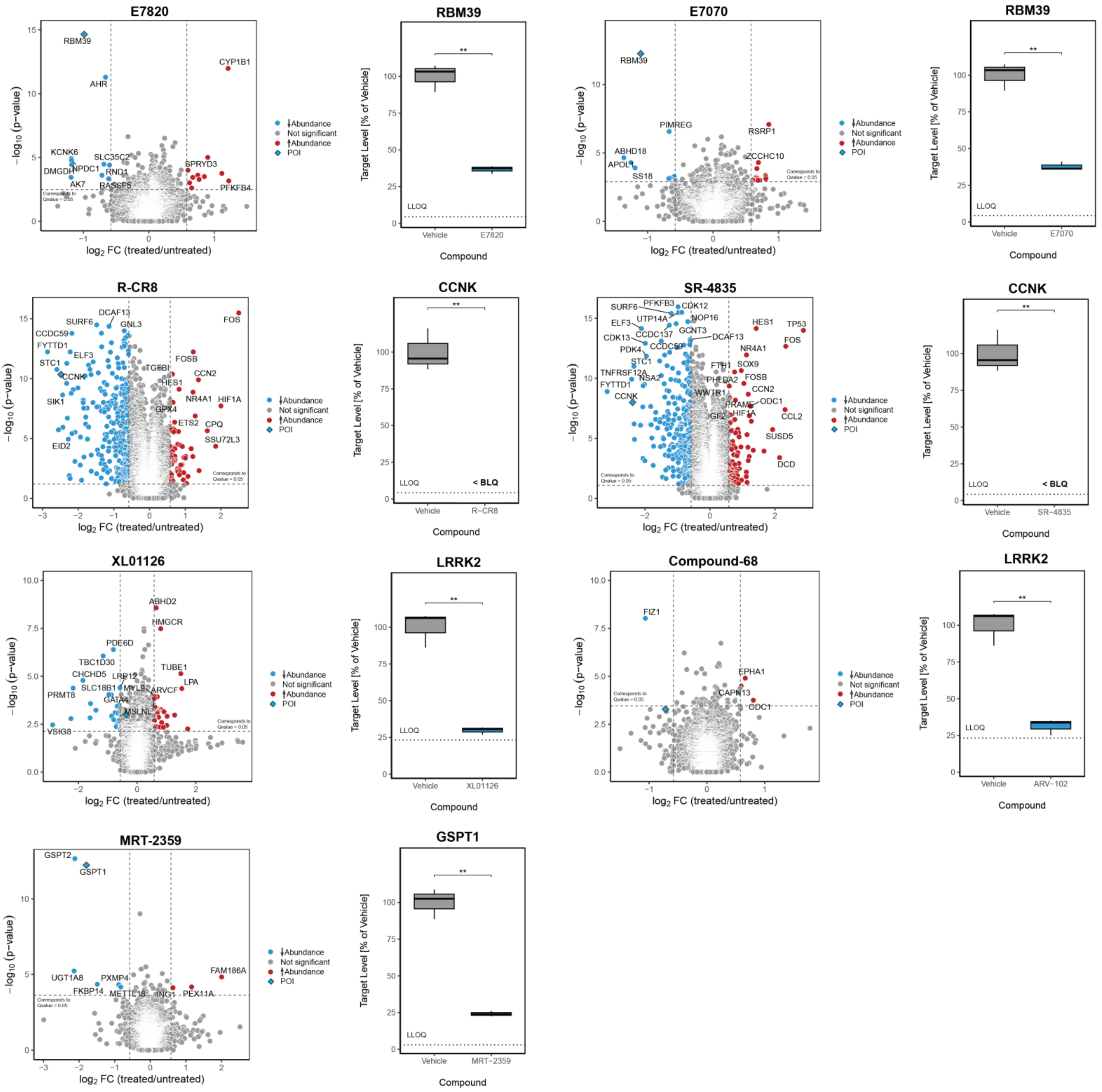


***Supplementary Figure 17 Automated sample preparation coupled to LC-MS can be utilized for target identification, quantification and selectivity profiling of molecular degraders.*** *Volcano plot illustrating the differentially abundant proteins identified through proteome profiling between A549 cell samples (n = 5) treated with the molecular degrader compounds (Supplementary Data 4, Supplementary Figure 17). Proteins are color-coded based on their statistical significance and fold change: red indicates increased abundance, blue indicates decreased abundance (|log2(FC)| > 0.58, q-value < 0.05) and grey indicates no significant change (|log2(FC)| ≤ 0.58, q-value > 0.05). Boxplot showing the absolute amount of the POIs in A549 cell samples (n = 3) treated with the molecular degrader compounds, compared to DMSO-treated samples (vehicle). T-test based p-values are indicated in the plots: * = p < 0.05, ** = p < 0.01, *** = p < 0.001, **** = p < 0.0001, n.s. = not significant. Lower limit of quantification (LLOQ) is indicated as dotted line. Targets falling below the lower limit of quantification (LLOQ) are marked as below limit of quantification (<BLQ).*

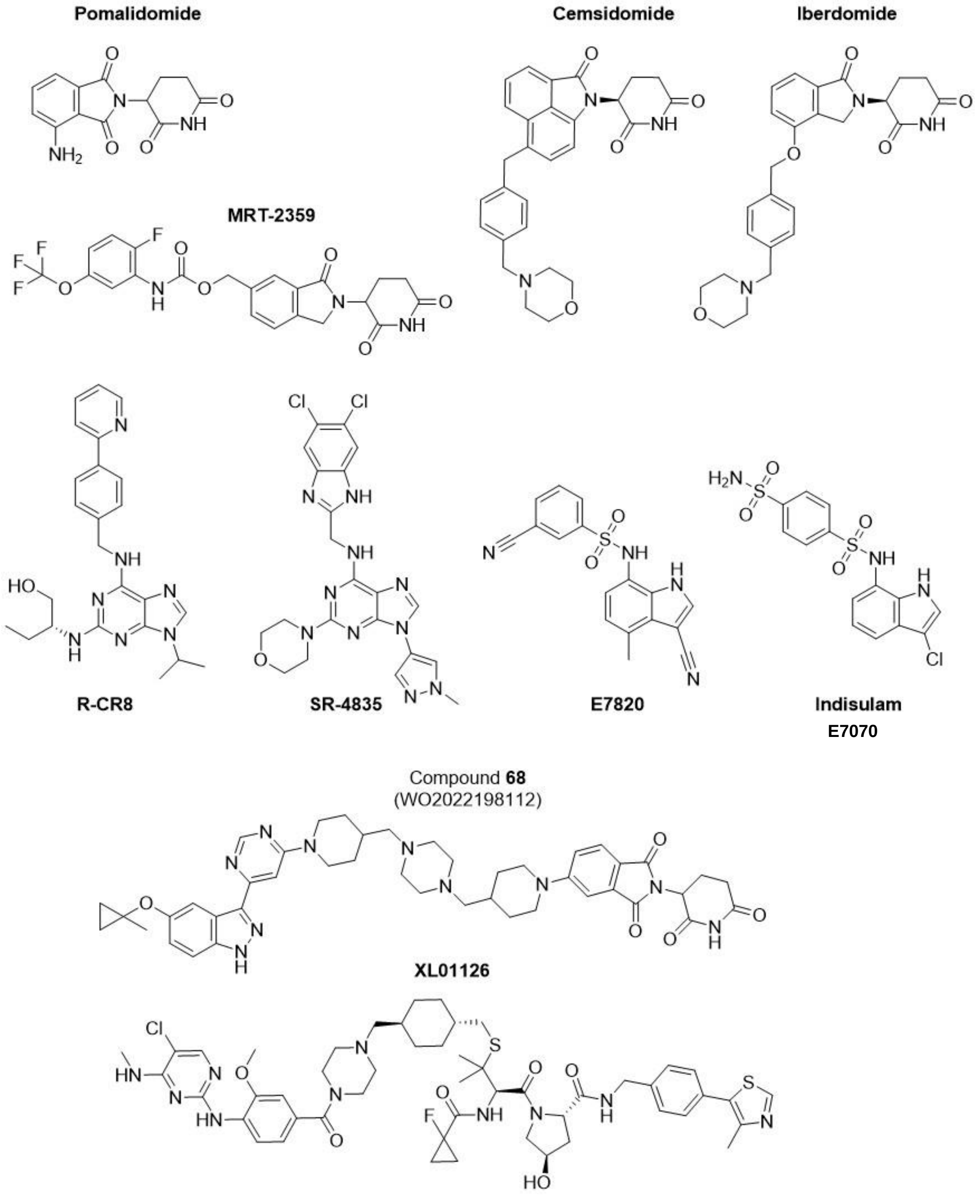


***Supplementary Figure 18 Automated sample preparation coupled to LC-MS can be utilized for target identification, quantification and selectivity profiling of molecular degraders.*** *Chemical structures of the tested preclinical and clinical-stage molecular degrader compounds (Supplementary Data 4).*

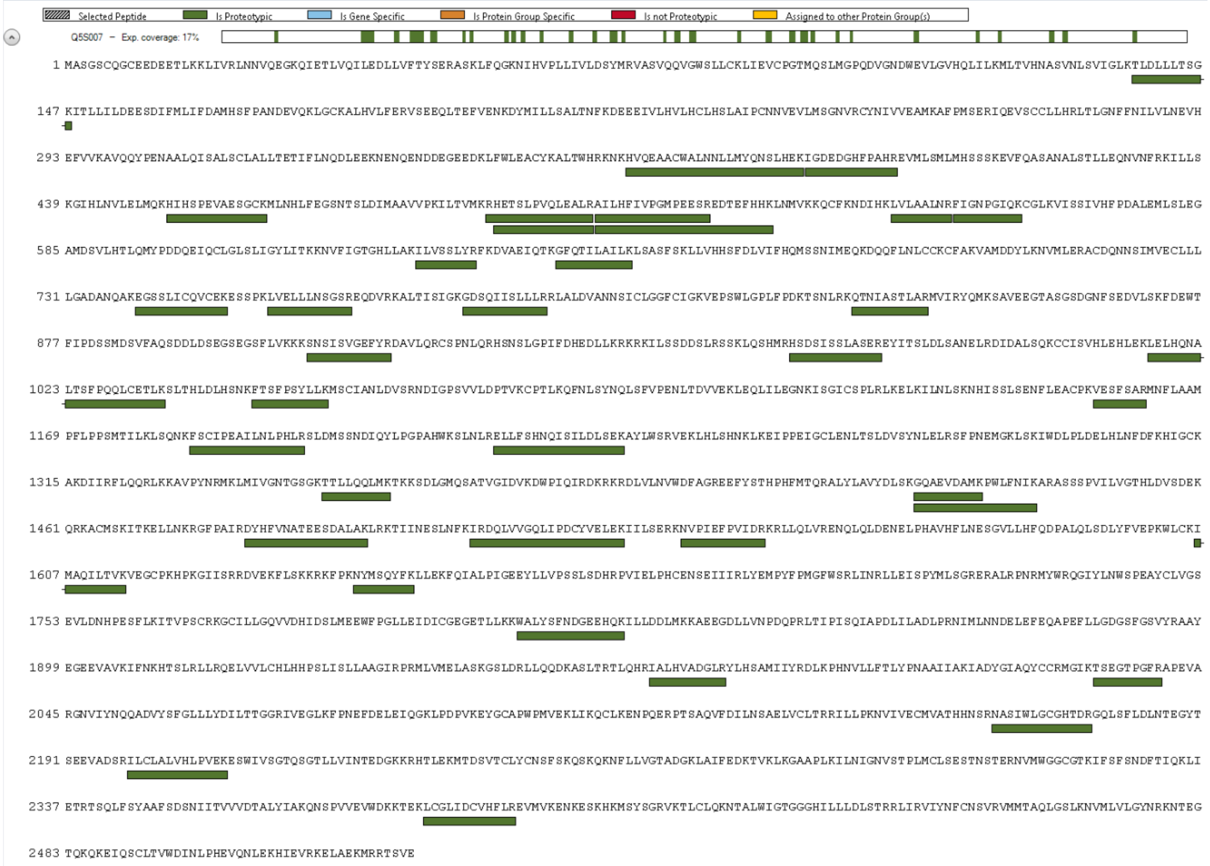


***Supplementary Figure 19 Sequence coverage of LRRK2 protein (UniProt ID - Q5S007)) in the proteome library after a proteomic profiling.*** *Exp. coverage % indicates the proportion of the full protein sequence identified. Green bars underline each proteotypic peptide of the protein that has been confidently identified in either A549 or K562 cell line.*

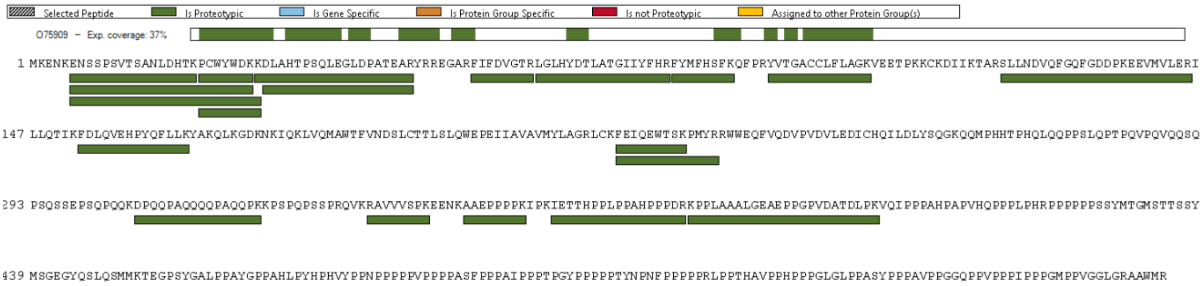


***Supplementary Figure 20 Sequence coverage of CCNK protein (UniProt ID – O75909)) in the proteome library after a proteomic profiling.*** *Exp. coverage % indicates the proportion of the full protein sequence identified. Green bars underline each proteotypic peptide of the protein that has been confidently identified in either A549 or K562 cell line.*

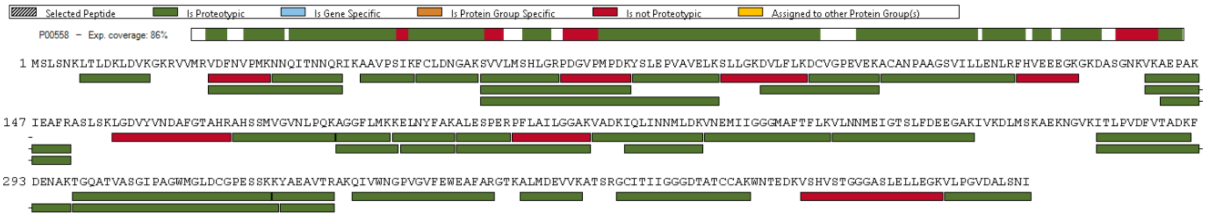


***Supplementary Figure 21 Sequence coverage of PGK1 protein (UniProt ID – P00558)) in the proteome library after a proteomic profiling.*** *Exp. coverage % indicates the proportion of the full protein sequence identified. Green bars underline each proteotypic peptide of the protein that has been confidently identified in either A549 or K562 cell line.*

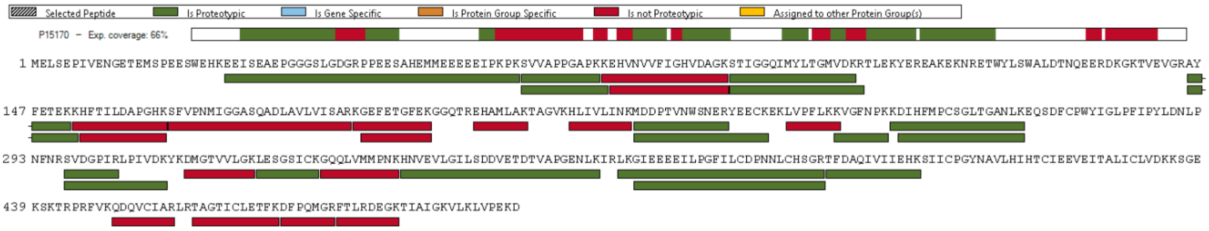


***Supplementary Figure 22 Sequence coverage of ERF3A protein (UniProt ID – P15170)) in the proteome library after a proteomic profiling.*** *Exp. coverage % indicates the proportion of the full protein sequence identified. Green bars underline each proteotypic peptide of the protein that has been confidently identified in either A549 or K562 cell line.*

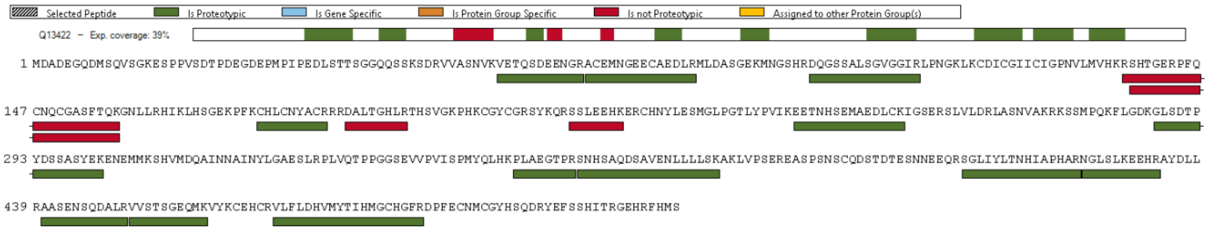


***Supplementary Figure 23 Sequence coverage of IKZF1 protein (UniProt ID – Q13422)) in the proteome library after a proteomic profiling.*** *Exp. coverage % indicates the proportion of the full protein sequence identified. Green bars underline each proteotypic peptide of the protein that has been confidently identified in either A549 or K562 cell line.*

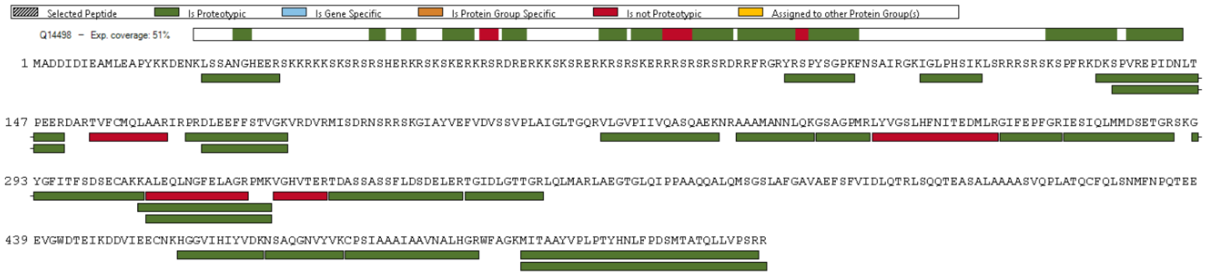


***Supplementary Figure 24 Sequence coverage of RBM39 protein (UniProt ID – Q14498)) in the proteome library after a proteomic profiling.*** *Exp. coverage % indicates the proportion of the full protein sequence identified. Green bars underline each proteotypic peptide of the protein that has been confidently identified in either A549 or K562 cell line.*

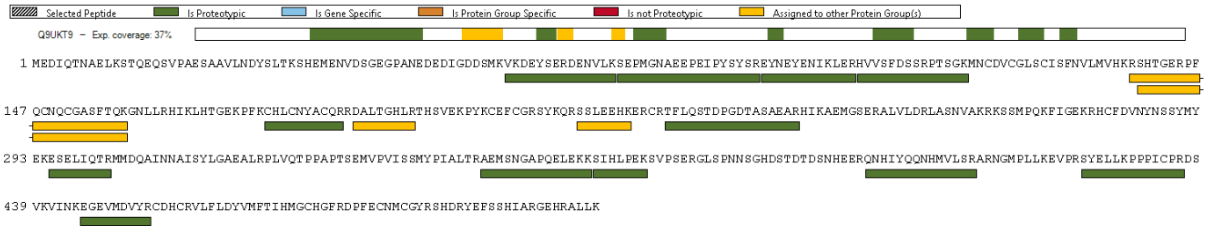


***Supplementary Figure 25 Sequence coverage of IKZF3 protein (UniProt ID – Q9UKT9)) in the proteome library after a proteomic profiling.*** *Exp. coverage % indicates the proportion of the full protein sequence identified. Green bars underline each proteotypic peptide of the protein that has been confidently identified in either A549 or K562 cell line.*

| | Assay |
|---|---|
| Accession | O75909 |
| Gene | CCNK |
| Protein Description | Cyclin − K |
| Peptide Stripped Sequence | FEIQEWTSK |
| Peptide Ion Variant | FEIQEWTSK (2+) |
| Calibrator Sequence | FEIQEWTSK[Lys8 C-term] |

#### Assay Characterization Results

| Parameter | Value | Unit |
|---|---|---|
| LLOQ | 0.0064 | fmol/ug |
| Average CV > LLOQ | 4.44 | % |
| Minimal Tested Concentration | 0.000256 | fmol/ug |
| Maximal Tested Concentration | 4 | fmol/ug |
| $R^2$ | 0.997 | |
| Slope (Linear Range) | 0.855 | |
| Intercept (Linear Range) | 8.01 | |

#### Assay Characterization Plot

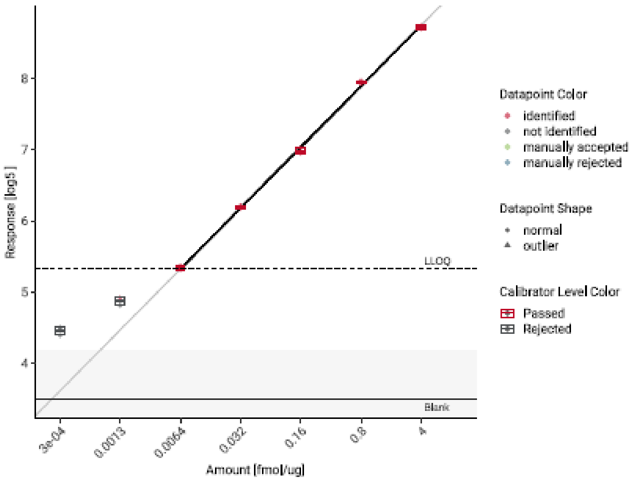


#### Assay Characterization Settings

| Parameter | Value |
|---|---|
| q-value Filter | < 0.05 |
| Zero Calibrator Filter | 3 x blank intensity |
| Accuracy Filter | <20 % Deviation |
| Precision Filter | < 20 % CV |

***Supplementary Figure 26 LLOQ determination of CCNK in K562.*** *7-point, 5 fold dilution curve of the best performing peptide of the protein CCNK was measured. LLOQ was determined after applying q-value filter of 0.05, Zero Calibrator Filter for 3x blank intensity, 20% accuracy filter, and 20% precision filter.*

| | Assay |
|---|---|
| Accession | P00558 |
| Gene | PGK1 |
| Protein Description | Phosphoglycerate kinase 1 |
| Peptide Stripped Sequence | YSLEPVAVELK |
| Peptide Ion Variant | YSLEPVAVELK (2+) |
| Calibrator Sequence | YSLEPVAVELK[Lys8 C-term] |

#### Assay Characterization Results

| Parameter | Value | Unit |
|---|---|---|
| LLOQ | 0.0064 | fmol/ug |
| Average CV > LLOQ | 6.22 | % |
| Minimal Tested Concentration | 0.000256 | fmol/ug |
| Maximal Tested Concentration | 4 | fmol/ug |
| $R^2$ | 0.993 | |
| Slope (Linear Range) | 0.946 | |
| Intercept (Linear Range) | 7.89 | |

#### Assay Characterization Plot

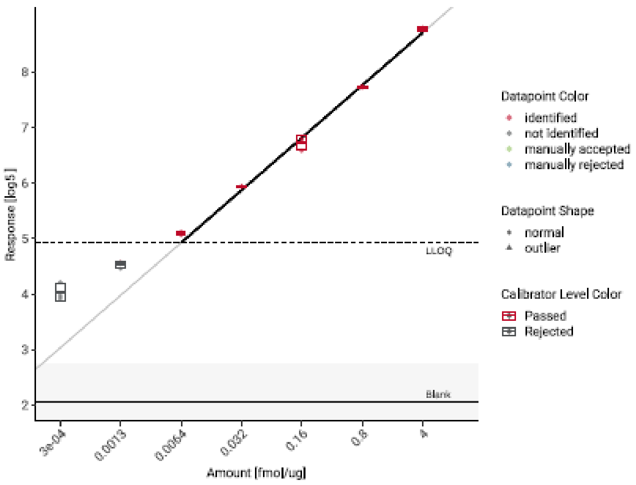


#### Assay Characterization Settings

| Parameter | Value |
|---|---|
| q-value Filter | < 0.05 |
| Zero Calibrator Filter | 3 x blank intensity |
| Accuracy Filter | <20 % Deviation |
| Precision Filter | < 20 % CV |

***Supplementary Figure 27 LLOQ determination of PGK1 in K562.*** *7-point, 5 fold dilution curve of the best performing peptide of the protein PGK1 was measured. LLOQ was determined after applying q-value filter of 0.05, Zero Calibrator Filter for 3x blank intensity, 20% accuracy filter, and 20% precision filter.*

| Assay | |
|---|---|
| Accession | P15170 |
| Gene | GSPT1 |
| Protein Description | Eukaryotic peptide chain release factor GTP-binding subunit ERF3A |
| Peptide Stripped Sequence | SVDGPIR |
| Peptide Ion Variant | SVDGPIR (2+) |
| Calibrator Sequence | SVDGPIR[Arg10 C-term] |

#### Assay Characterization Results

| Parameter | Value | Unit |
|---|---|---|
| LLOQ | 0.032 | fmol/ug |
| Average CV > LLOQ | 5.16 | % |
| Minimal Tested Concentration | 0.000256 | fmol/ug |
| Maximal Tested Concentration | 4 | fmol/ug |
| $R^2$ | 0.983 | |
| Slope (Linear Range) | 0.801 | |
| Intercept (Linear Range) | 8.07 | |

#### Assay Characterization Plot

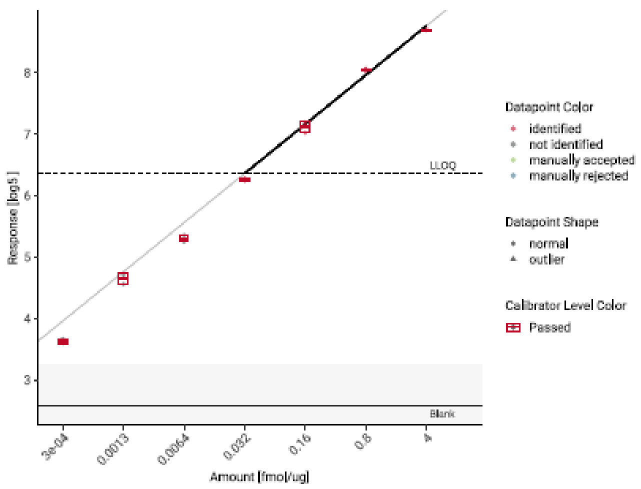


#### Assay Characterization Settings

| Parameter | Value |
|---|---|
| q-value Filter | < 0.05 |
| Zero Calibrator Filter | 3 x blank intensity |
| Accuracy Filter | <20 % Deviation |
| Precision Filter | < 20 % CV |

***Supplementary Figure 28 LLOQ determination of GSPT1 in K562.*** *7-point, 5 fold dilution curve of the best performing peptide of the protein GSPT1 was measured. LLOQ was determined after applying q-value filter of 0.05, Zero Calibrator Filter for 3x blank intensity, 20% accuracy filter, and 20% precision filter.*

| | Assay |
|---|---|
| Accession | Q13422 |
| Gene | IKZF1 |
| Protein Description | DNA-binding protein Ikaros |
| Peptide Stripped Sequence | SNHSAQDSAVENLLLLSK |
| Peptide Ion Variant | SNHSAQDSAVENLLLLSK (3+) |
| Calibrator Sequence | SNHSAQDSAVENLLLLSK[Lys8 C-term] |

#### Assay Characterization Results

| Parameter | Value | Unit |
|---|---|---|
| LLOQ | 0.00128 | fmol/ug |
| Average CV > LLOQ | 4.85 | % |
| Minimal Tested Concentration | 0.000256 | fmol/ug |
| Maximal Tested Concentration | 4 | fmol/ug |
| $R^2$ | 0.98 | |
| Slope (Linear Range) | 0.794 | |
| Intercept (Linear Range) | 7.92 | |

#### Assay Characterization Plot

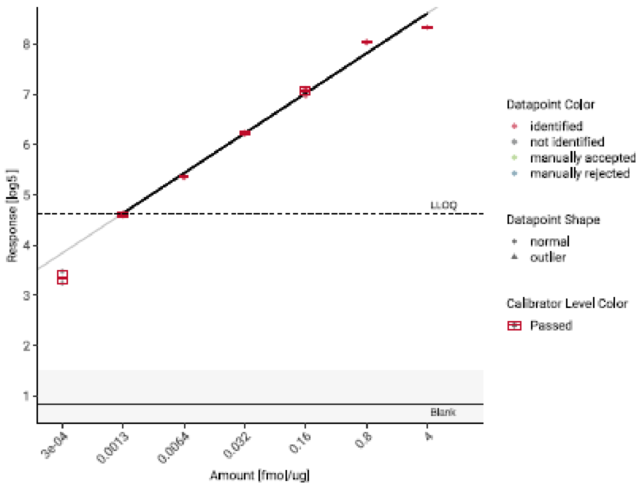


#### Assay Characterization Settings

| Parameter | Value |
|---|---|
| q-value Filter | < 0.05 |
| Zero Calibrator Filter | 3 x blank intensity |
| Accuracy Filter | <20 % Deviation |
| Precision Filter | < 20 % CV |

***Supplementary Figure 29 LLOQ determination of IKZF1 in K562.*** *7-point, 5 fold dilution curve of the best performing peptide of the protein IKZF1 was measured. LLOQ was determined after applying q-value filter of 0.05, Zero Calibrator Filter for 3x blank intensity, 20% accuracy filter, and 20% precision filter.*

| | Assay |
|---|---|
| Accession | Q14498 |
| Gene | RBM39 |
| Protein Description | RNA-binding protein 39 |
| Peptide Stripped Sequence | TGIDLGTTGR |
| Peptide Ion Variant | TGIDLGTTGR (2+) |
| Calibrator Sequence | TGIDLGTTGR[Arg10 C-term] |

#### Assay Characterization Results

| Parameter | Value | Unit |
|---|---|---|
| LLOQ | 0.032 | fmol/ug |
| Average CV > LLOQ | 7.15 | % |
| Minimal Tested Concentration | 0.000256 | fmol/ug |
| Maximal Tested Concentration | 4 | fmol/ug |
| $R^2$ | 0.995 | |
| Slope (Linear Range) | 0.976 | |
| Intercept (Linear Range) | 8.4 | |

#### Assay Characterization Plot

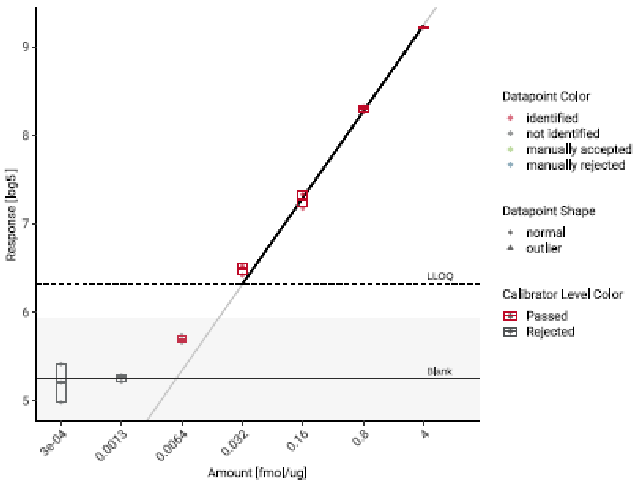


#### Assay Characterization Settings

| Parameter | Value |
|---|---|
| q-value Filter | < 0.05 |
| Zero Calibrator Filter | 3 x blank intensity |
| Accuracy Filter | <20 % Deviation |
| Precision Filter | < 20 % CV |

***Supplementary Figure 30 LLOQ determination of RBM39 in K562.*** *7-point, 5 fold dilution curve of the best performing peptide of the protein RBM39 was measured. LLOQ was determined after applying q-value filter of 0.05, Zero Calibrator Filter for 3x blank intensity, 20% accuracy filter, and 20% precision filter.*

| Assay | |
|---|---|
| Accession | Q5S007 |
| Gene | LRRK2 |
| Protein Description | Leucine-rich repeat serine/threonine-protein kinase 2 |
| Peptide Stripped Sequence | VESFSAR |
| Peptide Ion Variant | VESFSAR (2+) |
| Calibrator Sequence | VESFSAR[Arg10 C-term] |

Assay Characterization Results

| Parameter | Value | Unit |
|---|---|---|
| LLOQ | 0.032 | fmol/ug |
| Average CV > LLOQ | 4.28 | % |
| Minimal Tested Concentration | 0.000256 | fmol/ug |
| Maximal Tested Concentration | 4 | fmol/ug |
| $R^2$ | 0.998 | |
| Slope (Linear Range) | 0.986 | |
| Intercept (Linear Range) | 7.98 | |

Assay Characterization Settings

| Parameter | Value |
|---|---|
| q-value Filter | < 0.05 |
| Zero Calibrator Filter | 3 x blank intensity |
| Accuracy Filter | <20 % Deviation |
| Precision Filter | < 20 % CV |

Assay Characterization Plot

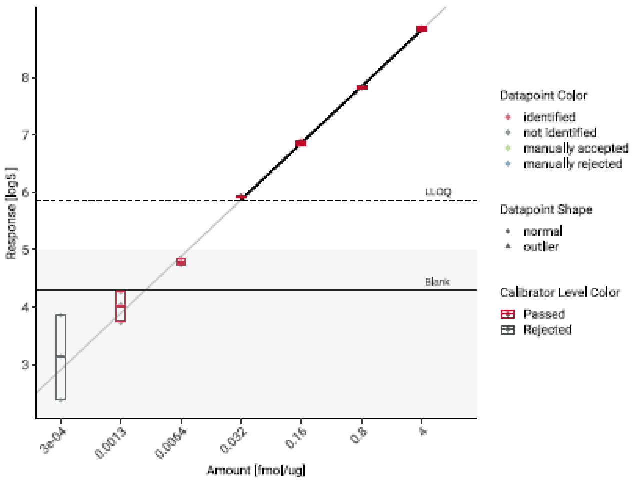


***Supplementary Figure 31 LLOQ determination of LRRK2 in K562.*** *7-point, 5 fold dilution curve of the best performing peptide of the protein LRRK2 was measured. LLOQ was determined after applying q-value filter of 0.05, Zero Calibrator Filter for 3x blank intensity, 20% accuracy filter, and 20% precision filter.*

| | Assay |
|---|---|
| Accession | Q9UKT9 |
| Gene | IKZF3 |
| Protein Description | Zinc finger protein Aiolos |
| Peptide Stripped Sequence | SIHLPEK |
| Peptide Ion Variant | SIHLPEK (2+) |
| Calibrator Sequence | SIHLPEK[Lys8 C-term] |

#### Assay Characterization Results

| Parameter | Value | Unit |
|---|---|---|
| LLOQ | 0.032 | fmol/ug |
| Average CV > LLOQ | 3.05 | % |
| Minimal Tested Concentration | 0.000256 | fmol/ug |
| Maximal Tested Concentration | 4 | fmol/ug |
| $R^2$ | 0.998 | |
| Slope (Linear Range) | 0.892 | |
| Intercept (Linear Range) | 7.09 | |

#### Assay Characterization Settings

| Parameter | Value |
|---|---|
| q-value Filter | < 0.05 |
| Zero Calibrator Filter | 3 x blank intensity |
| Accuracy Filter | <20 % Deviation |
| Precision Filter | < 20 % CV |

#### Assay Characterization Plot

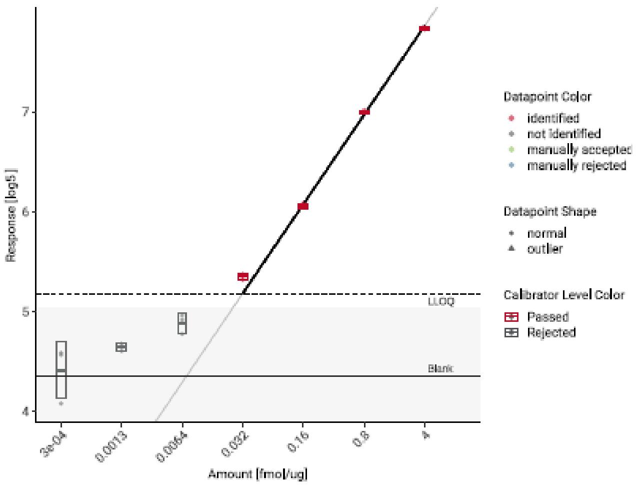


***Supplementary Figure 32 LLOQ determination of IKZF3 in K562.*** *7-point, 5 fold dilution curve of the best performing peptide of the protein IKZF3 was measured. LLOQ was determined after applying q-value filter of 0.05, Zero Calibrator Filter for 3x blank intensity, 20% accuracy filter, and 20% precision filter.*

| Assay | |
|---|---|
| Accession | P15170 |
| Gene | GSPT1 |
| Protein Description | Eukaryotic peptide chain release factor GTP-binding subunit ERF3A |
| Peptide Stripped Sequence | SVDGPIR |
| Peptide Ion Variant | SVDGPIR (2+) |
| Calibrator Sequence | SVDGPIR[Arg10 C-term] |

#### Assay Characterization Results

| Parameter | Value | Unit |
|---|---|---|
| LLOQ | 0.16 | fmol/ug |
| Average CV > LLOQ | 5.36 | % |
| Minimal Tested Concentration | 0.000256 | fmol/ug |
| Maximal Tested Concentration | 4 | fmol/ug |
| $R^2$ | 0.996 | |
| Slope (Linear Range) | 0.904 | |
| Intercept (Linear Range) | 7.9 | |

#### Assay Characterization Plot

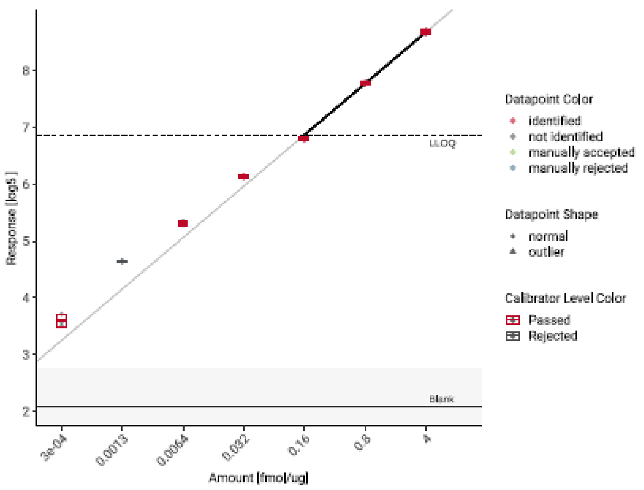


#### Assay Characterization Settings

| Parameter | Value |
|---|---|
| q-value Filter | < 0.05 |
| Zero Calibrator Filter | 3 x blank intensity |
| Accuracy Filter | <20 % Deviation |
| Precision Filter | < 20 % CV |

***Supplementary Figure 33 LLOQ determination of GSPT1 in A549.*** *7-point, 5 fold dilution curve of the best performing peptide of the protein GSPT1 was measured. LLOQ was determined after applying q-value filter of 0.05, Zero Calibrator Filter for 3x blank intensity, 20% accuracy filter, and 20% precision filter.*

| | Assay |
|---|---|
| Accession | O75909 |
| Gene | CCNK |
| Protein Description | Cyclin – K |
| Peptide Stripped Sequence | FEIQEWTSK |
| Peptide Ion Variant | FEIQEWTSK (2+) |
| Calibrator Sequence | FEIQEWTSK[Lys8 C-term] |

Assay Characterization Results

| Parameter | Value | Unit |
|---|---|---|
| LLOQ | 0.0064 | fmol/ug |
| Average CV > LLOQ | 3.83 | % |
| Minimal Tested Concentration | 0.000256 | fmol/ug |
| Maximal Tested Concentration | 4 | fmol/ug |
| $R^2$ | 0.998 | |
| Slope (Linear Range) | 0.868 | |
| Intercept (Linear Range) | 7.87 | |

Assay Characterization Settings

| Parameter | Value |
|---|---|
| q-value Filter | < 0.05 |
| Zero Calibrator Filter | 3 x blank intensity |
| Accuracy Filter | <20 % Deviation |
| Precision Filter | < 20 % CV |

Assay Characterization Plot

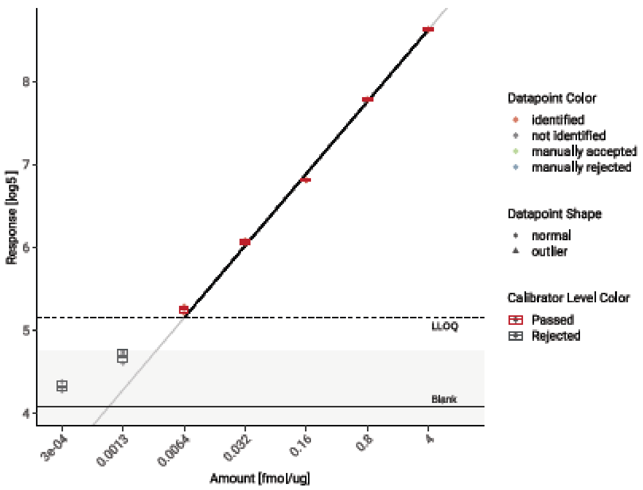


***Supplementary Figure 34 LLOQ determination of CCNK in A549.*** *7-point, 5 fold dilution curve of the best performing peptide of the protein CCNK was measured. LLOQ was determined after applying q-value filter of 0.05, Zero Calibrator Filter for 3x blank intensity, 20% accuracy filter, and 20% precision filter.*

| | Assay |
|---|---|
| Accession | Q5S007 |
| Gene | LRRK2 |
| Protein Description | Leucine-rich repeat serine/threonine-protein kinase 2 |
| Peptide Stripped Sequence | VESFSAR |
| Peptide Ion Variant | VESFSAR (2+) |
| Calibrator Sequence | VESFSAR[Arg10 C-term] |

Assay Characterization Results

| Parameter | Value | Unit |
|---|---|---|
| LLOQ | 0.0064 | fmol/ug |
| Average CV > LLOQ | 10.2 | % |
| Minimal Tested Concentration | 0.000256 | fmol/ug |
| Maximal Tested Concentration | 4 | fmol/ug |
| $R^2$ | 0.989 | |
| Slope (Linear Range) | 0.904 | |
| Intercept (Linear Range) | 7.4 | |

Assay Characterization Settings

| Parameter | Value |
|---|---|
| q-value Filter | < 0.05 |
| Zero Calibrator Filter | 3 x blank intensity |
| Accuracy Filter | <20 % Deviation |
| Precision Filter | < 20 % CV |

Assay Characterization Plot

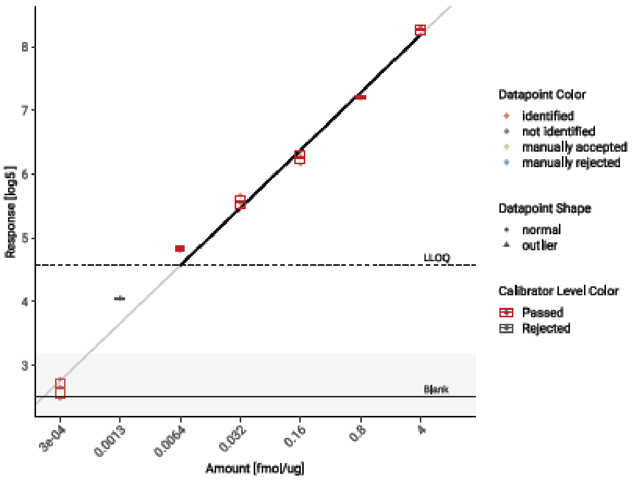


***Supplementary Figure 35 LLOQ determination of LRRK2 in A549.*** *7-point, 5 fold dilution curve of the best performing peptide of the protein LRRK2 was measured. LLOQ was determined after applying q-value filter of 0.05, Zero Calibrator Filter for 3x blank intensity, 20% accuracy filter, and 20% precision filter.*

| | Assay |
|---|---|
| Accession | Q14498 |
| Gene | RBM39 |
| Protein Description | RNA-binding protein 39 |
| Peptide Stripped Sequence | TGIDLGTTGR |
| Peptide Ion Variant | TGIDLGTTGR (2+) |
| Calibrator Sequence | TGIDLGTTGR[Arg10 C-term] |

Assay Characterization Results

| Parameter | Value | Unit |
|---|---|---|
| LLOQ | 0.16 | fmol/ug |
| Average CV > LLOQ | 4.82 | % |
| Minimal Tested Concentration | 0.000256 | fmol/ug |
| Maximal Tested Concentration | 4 | fmol/ug |
| $R^2$ | 0.997 | |
| Slope (Linear Range) | 1.06 | |
| Intercept (Linear Range) | 8.3 | |

Assay Characterization Plot

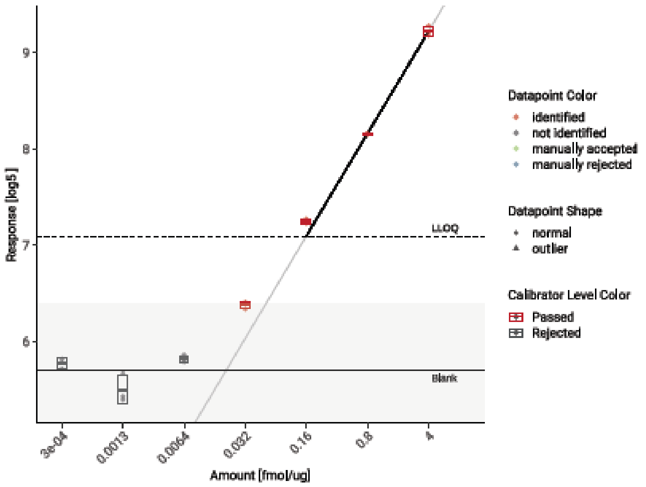


Assay Characterization Settings

| Parameter | Value |
|---|---|
| q-value Filter | < 0.05 |
| Zero Calibrator Filter | 3 x blank intensity |
| Accuracy Filter | <20 % Deviation |
| Precision Filter | < 20 % CV |

***Supplementary Figure 36 LLOQ determination of RBM39 in A549.*** *7-point, 5 fold dilution curve of the best performing peptide of the protein RBM39 was measured. LLOQ was determined after applying q-value filter of 0.05, Zero Calibrator Filter for 3x blank intensity, 20% accuracy filter, and 20% precision filter.*

| | Assay |
|---|---|
| Accession | P00558 |
| Gene | PGK1 |
| Protein Description | Phosphoglycerate kinase 1 |
| Peptide Stripped Sequence | YSLEPVAVELK |
| Peptide Ion Variant | YSLEPVAVELK (2+) |
| Calibrator Sequence | YSLEPVAVELK[Lys8 C-term] |

Assay Characterization Results

| Parameter | Value | Unit |
|---|---|---|
| LLOQ | 0.032 | fmol/ug |
| Average CV > LLOQ | 2.76 | % |
| Minimal Tested Concentration | 0.000256 | fmol/ug |
| Maximal Tested Concentration | 4 | fmol/ug |
| $R^2$ | 0.998 | |
| Slope (Linear Range) | 1.01 | |
| Intercept (Linear Range) | 7.82 | |

Assay Characterization Plot

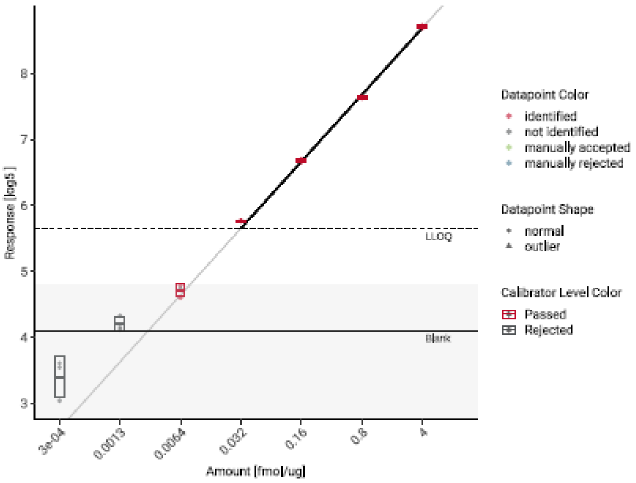


Assay Characterization Settings

| Parameter | Value |
|---|---|
| q-value Filter | < 0.05 |
| Zero Calibrator Filter | 3 x blank intensity |
| Accuracy Filter | <20 % Deviation |
| Precision Filter | < 20 % CV |

***Supplementary Figure 37 LLOQ determination of PGK1 in A549.*** *7-point, 5 fold dilution curve of the best performing peptide of the protein PGK1 was measured. LLOQ was determined after applying q-value filter of 0.05, Zero Calibrator Filter for 3x blank intensity, 20% accuracy filter, and 20% precision filter.*

| | Assay |
|---|---|
| Accession | Q9UKT9 |
| Gene | IKZF3 |
| Protein Description | Zinc finger protein Aiolos |
| Peptide Stripped Sequence | SIHLPEK |
| Peptide Ion Variant | SIHLPEK (2+) |
| Calibrator Sequence | SIHLPEK[Lys8 C-term] |

#### Assay Characterization Results

| Parameter | Value | Unit |
|---|---|---|
| LLOQ | 0.16 | fmol/ug |
| Average CV > LLOQ | 3.82 | % |
| Minimal Tested Concentration | 0.000256 | fmol/ug |
| Maximal Tested Concentration | 4 | fmol/ug |
| $R^2$ | 0.998 | |
| Slope (Linear Range) | 0.985 | |
| Intercept (Linear Range) | 7.26 | |

#### Assay Characterization Plot

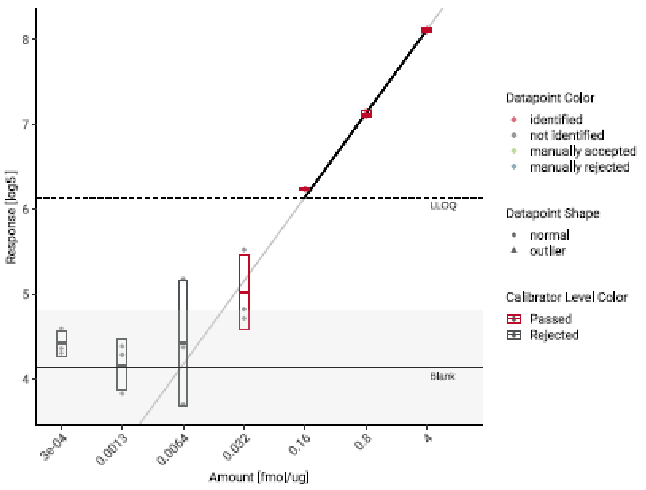


#### Assay Characterization Settings

| Parameter | Value |
|---|---|
| q-value Filter | < 0.05 |
| Zero Calibrator Filter | 3 x blank intensity |
| Accuracy Filter | <20 % Deviation |
| Precision Filter | < 20 % CV |

***Supplementary Figure 38 LLOQ determination of IKZF3 in A549.*** *7-point, 5 fold dilution curve of the best performing peptide of the protein IKZF3 was measured. LLOQ was determined after applying q-value filter of 0.05, Zero Calibrator Filter for 3x blank intensity, 20% accuracy filter, and 20% precision filter.*

| | Assay |
|---|---|
| Accession | Q13422 |
| Gene | IKZF1 |
| Protein Description | DNA-binding protein Ikaros |
| Peptide Stripped Sequence | SNHSAQDSAVENLLLLSK |
| Peptide Ion Variant | SNHSAQDSAVENLLLLSK (3+) |
| Calibrator Sequence | SNHSAQDSAVENLLLLSK[Lys8 C-term] |

#### Assay Characterization Results

| Parameter | Value | Unit |
|---|---|---|
| LLOQ | 0.0064 | fmol/ug |
| Average CV > LLOQ | 5.7 | % |
| Minimal Tested Concentration | 0.000256 | fmol/ug |
| Maximal Tested Concentration | 4 | fmol/ug |
| $R^2$ | 0.987 | |
| Slope (Linear Range) | 0.792 | |
| Intercept (Linear Range) | 7.7 | |

#### Assay Characterization Plot

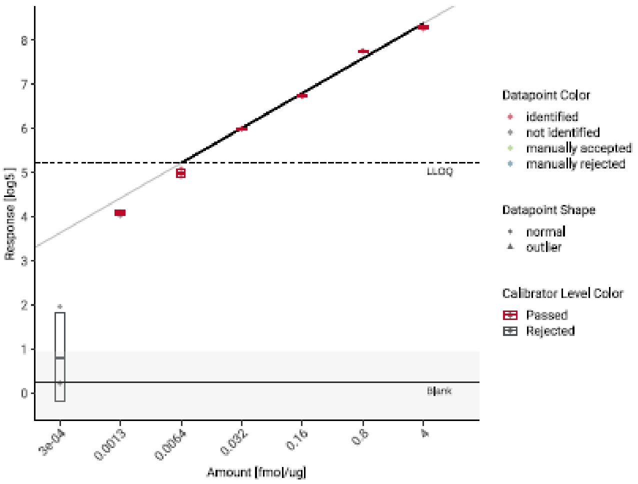


#### Assay Characterization Settings

| Parameter | Value |
|---|---|
| q-value Filter | < 0.05 |
| Zero Calibrator Filter | 3 x blank intensity |
| Accuracy Filter | <20 % Deviation |
| Precision Filter | < 20 % CV |

***Supplementary Figure 39 LLOQ determination of IKZF1 in A549.*** *7-point, 5 fold dilution curve of the best performing peptide of the protein IKZF1 was measured. LLOQ was determined after applying q-value filter of 0.05, Zero Calibrator Filter for 3x blank intensity, 20% accuracy filter, and 20% precision filter.*

#### References

1. Fu, T. J., Peng, J., Lee, G., Price, D. H. & Flores, O. Cyclin K functions as a CDK9 regulatory subunit and participates in RNA polymerase II transcription. *J. Biol. Chem.* **274**, 34527–34530 (1999).
2. Houles, T. *et al.* The CDK12 inhibitor SR-4835 functions as a molecular glue that promotes cyclin K degradation in melanoma. *Cell Death Discov.* **9**, 459 (2023).
3. Słabicki, M. *et al.* The CDK inhibitor CR8 acts as a molecular glue degrader that depletes cyclin K. *Nature* **585**, 293–297 (2020).
4. Law, J. C., Ritke, M. K., Yalowich, J., Leder, G. & Ferrell, R. Mutational inactivation of the p53 gene in the human erythroid leukemic K562 cell line. *Leuk. Res.* **17**, 1045–1050 (1993).
